# Dynamic accumulation of a helper NLR at the plant-pathogen interface underpins pathogen recognition

**DOI:** 10.1101/2021.03.15.435521

**Authors:** Cian Duggan, Eleonora Moratto, Zachary Savage, Eranthika Hamilton, Hiroaki Adachi, Chih-Hang Wu, Alexandre Y Leary, Yasin Tumtas, Stephen M Rothery, Abbas Maqbool, Seda Nohut, Sophien Kamoun, Tolga Osman Bozkurt

## Abstract

Plants employ sensor-helper pairs of NLR immune receptors to recognize pathogen effectors and activate immune responses. Yet the subcellular localization of NLRs pre- and post-activation during pathogen infection remains poorly known. Here we show that NRC4, from the ‘NRC’ solanaceous helper NLR family, undergoes dynamic changes in subcellular localization by shuttling to and from the plant-pathogen haustorium interface established during infection by the Irish potato famine pathogen *Phytophthora infestans.* Specifically, prior to activation, NRC4 accumulates at the extra-haustorial membrane (EHM), presumably to mediate response to perihaustorial effectors, that are recognized by NRC4-dependent sensor NLRs. However not all NLRs accumulate at the EHM, as the closely related helper NRC2, and the distantly related ZAR1, did not accumulate at the EHM. NRC4 required an intact N-terminal coiled coil domain to accumulate at the EHM, whereas the functionally conserved MADA motif implicated in cell death activation and membrane insertion was dispensable for this process. Strikingly, a constitutively autoactive NRC4 mutant did not accumulate at the EHM and showed punctate distribution that mainly associated with the plasma membrane, suggesting that post-activation, NRC4 probably undergoes a conformation switch to form clusters that do not preferentially associate with the EHM. When NRC4 is activated by a sensor NLR during infection however, NRC4 formed puncta mainly at the EHM and to a lesser extent at the plasma membrane. We conclude that following activation at the EHM, NRC4 may spread to other cellular membranes from its primary site of activation to trigger immune responses.

**Significance statement:** Plant NLR immune receptors function as intracellular sensors of pathogen virulence factors known as effectors. In resting state, NLRs localize to subcellular sites where the effectors they sense operate. However, the extent to which NLRs alter their subcellular distribution during infection remains elusive. We describe dynamic changes in spatiotemporal localization of an NLR protein in infected plant cells. Specifically, the NLR protein accumulates at the newly synthesized plant-pathogen interface membrane, where the corresponding effectors are deployed. Following immune recognition, the activated receptor re-organizes to form punctate structures that target the cell periphery. We propose that NLRs are not necessarily stationary immune receptors, but instead may spread to other cellular membranes from the primary site of activation to boost immune responses.

## Introduction

Filamentous pathogens cause devastating diseases on crops, posing a major threat to food security. Some oomycete and fungal pathogens produce specialised hyphal extensions called haustoria that invade the host cells. Haustoria are critical infection structures implicated in delivery of effector proteins and nutrient uptake (1–6). These specialized infection structures are accommodated within the plant cells but are excluded from the host cytoplasm through a newly synthesized membrane called the extrahaustorial membrane (EHM). An intriguing, yet poorly understood, observation is that the EHM is continuous with the host plasma membrane but is distinct in lipid and protein composition (7, 8). Most of the proteins embedded in the plasma membrane, such as surface immune receptors, are excluded from the EHM (9–11). Despite its critical role as the ultimate interface mediating macromolecule exchange between the host and parasite, the mechanisms underlying the biogenesis and functions of the EHM are poorly understood (12).

Pathogens deliver effector proteins inside the host cells to neutralize immune responses and enable parasitic infection. A well-studied class of effectors delivered via haustorium are the RXLR family of effectors secreted by *P. infestans* (2, 6, 13). RXLR effectors traffic to diverse plant cell compartments to suppress host immunity and mediate nutrient uptake. Remarkably, several *P. infestans* RXLR effectors focally accumulate at the haustorium interface and perturb cellular defences (7, 13–15). These include AVRblb2 and AVR1, both of which are implicated in targeting host defense-related secretory pathways to contribute to pathogen virulence (14, 16). Notably, all *P. infestans* isolates harbour multiple *Avrblb2* paralogues (17), which are recognized by the broad-spectrum disease resistance gene *Rpi-blb2* cloned from the wild potato species *Solanum bulbocastanum* (14, 18, 19). On the other hand AVR1 is sensed by the late blight resistance gene *R1* which provides race specific resistance to AVR1 carrying *P. infestans* strains (20).

Both *Rpi-blb2* and *R1* encode nucleotide-binding, leucine-rich repeat proteins (NLR) which belong to an NLR network in Solanaceae and other asterid plants known as the NLR REQUIRED FOR CELL DEATH (NRC) family. The NRC immune network members form a superclade that consists of about one-third of all Solanaceae NLRs, providing disease resistance to nematodes, viruses, bacteria, oomycetes and aphids (21). Within the NRC network, sensor NLRs specialized to recognize effectors secreted by the pathogens are coupled to helper NLRs (NRCs) that translate the defense signal into disease resistance. We recently showed that Rpi-blb2 and R1 are ‘sensor NLRs’ that require the ‘helper NLR’ NRC4 for the immune-related programmed cell death known as the hypersensitive response (HR) and subsequent disease resistance (21). How and where AVRblb2 and AVR1 are recognized by the NRC4 helper-sensor pairs, as well as the mechanism that leads to HR and disease resistance following their recognition, are unknown. Because AVRblb2 and AVR1 localize to the EHM (13, 14), it is likely that the NLR receptor pairs that sense these perihaustorial effectors also accumulate at the haustorial interface. So far, live cell imaging of NLRs during infection, which would allow for a greater understanding of NLR functions, has not been feasible due to cell death activation. However, recently solved structures of activated NLRs uncovered critical residues that can be mutated to avoid HR activation without perturbing other NLR functions such as effector recognition and self-oligomerization following activation (22–24). Therefore, fluorescent protein fusions of these NLR mutants could be used for cell biology studies to investigate NLR activities during infection.

All NLRs in the NRC superclade carry N terminal coiled-coil (CC) domains, a characteristic of the CC-NLR type of immune receptors. The recently resolved cryogenic electron microscopy (Cryo-EM) structures of activated/non-activated forms of the CC-NLR type of resistance protein AtZAR1 (22, 25), which provides resistance to several bacterial species, revealed an intriguing model for HR elicitation. Upon activation, AtZAR1 (hereafter ZAR1) oligomerises into an inflammasome-like structure, called a ‘resistosome’. The ZAR1 resistosome consists of five ZAR1 proteins that assemble into a pentameric structure together with the kinases required for ZAR1 activation. Intriguingly, upon immune activation, the first alpha-helix (α1) within the CC domain of ZAR1 is exposed and the five α1 helices of the ZAR1 pentamer assemble into a funnel shaped structure. The resistosome is presumed to insert into the plasma membrane, forming a pore that could disrupt the cellular integrity or lead to ion flux across the membrane leading to an HR (26). This challenged the long-held view that NLRs execute HR and resistance through activation of downstream signalling cascades. However, the experimental evidence that supports this pore-forming activity and whether this model could be applied to other NLRs are still lacking. Furthermore, which cellular membrane the ZAR1 resistosome associates with remains to be determined.

We recently made an exciting discovery that an N-terminal motif (“the MADA motif”) overlapping ZAR1’s α1 helix, with the consensus sequence MADAxVSFxVxKLxxLLxxEx exists in ∼20% of CC-NLRs from monocot and dicot species. Remarkably, this motif is preserved in all NRC helpers but not in their sensor mates. Intriguingly, the first 29 amino acids of NRC4 containing the MADA motif elicited HR when fused to YFP on its C-terminus, but not when it is tagged with the YFP mutant that cannot oligomerize. These results indicated that the N-terminus of NRC4 relies on a scaffold such as YFP or the rest of NRC4, to form oligomers and trigger cell death. Notably, we previously showed that a chimeric NRC4 construct carrying ZAR1’s α1 helix is functional for triggering HR and confers disease resistance when co-expressed with Rpi-blb2 in lines lacking NRC4 (23), indicating that the proposed ZAR1 mode of action could be applied to NRC helpers.

Although recent structural studies greatly improved our understanding of the NLR mediated immunity and provide unprecedented insights into NLR mode of action, subcellular distribution of NLRs during infection with relevant pathogens is unknown. In the absence of infection, NLRs have been shown to localize to various cellular compartments such as cytoplasm, plasma membrane, nucleus and tonoplast etc (27–31). However, determining the subcellular localization of NLRs during infection has not been feasible due to activation of HR when the corresponding effector is present. Nevertheless, it is possible to monitor the distribution of the NRC helpers in plants that lack the sensor NLRs specialized to recognize the pathogen. The solanaceous model plant *Nicotiana benthamiana* is an excellent system to study the functioning of the Rpi-blb2-NRC4 pair as it contains a functional NRC4 but lacks specialized sensor NLRs that can recognize *P. infestans*, whereas transgenic plants carrying Rpi-blb2 are fully resistant to *P. infestans* (14, 18, 19)

Here we describe the dynamic changes in spatiotemporal localization of NRC4 in response to infection by *P. infestans.* NRC4 accumulates at the newly synthesized EHM, where the corresponding effectors AVRblb2 and AVR1 are deployed (13, 14). Following immune recognition, the activated receptor re-organizes to form punctate structures that target the cell periphery. Our results indicate that NLRs are not necessarily stationary immune receptors, but instead can alter their localisations during infection and may further spread to other cellular membranes from the primary site of activation to boost immune responses.

## Results

### Unlike NRC2 and ZAR1, non-activated NRC4 accumulates at the EHM during *Phytophthora infestans* infection

It has not been possible to investigate NLR subcellular localisation during infection with relevant microbes due to HR cell death. However, it is feasible to monitor helper NLRs during infection in the absence of sensor NLRs. *N. benthamiana-P. infestans* pathosystem offers excellent tools to investigate helper NLR functions because *N. benthamiana* lacks sensor NLRs that can prevent infection by *P. infestans*. *N. benthamiana* contains two alleles of NRC4 helper (NRC4a/b) which can pair with the sensor NLR Rpi-blb2 or R1 to recognize *P. infestans* effector protein AVRblb2 and R1 (14, 18, 19, 21, 23). Since both AVRblb2 and AVR1 accumulate at the haustorium interface (7, 13, 14), we reasoned that the corresponding sensors-helper pairs may be positioned at the EHM to detect these effectors.

Unfortunately, the sensor NLRs Rpi-blb2 and R1 produced too low fluorescence to accurately monitor during infection, and produced cell death under endogenous conditions – i.e. when NRC4 was present. Thus, we decided to investigate the localization of fluorescently tagged helper NRC4 in *N. benthamiana* during *P. infestans* infection. Strikingly, consistent with the localization pattern of the perihaustorial AVR effectors, both N-and C-terminal fusions of NRC4 accumulated around the haustorium when transiently expressed (Fig 1A & B). GFP fusions of NRC4 produced bright fluorescent signal around the haustorium but not throughout the cell, whereas in uninfected cells NRC4 showed mainly a cytoplasmic distribution much like the GFP control, except that NRC4 was excluded from the nucleus (Fig 1C & E-G). We then confirmed accumulation of NRC4 around the haustorium by infection microscopy assays in stable transgenic *35S::NRC4-GFP N. benthamiana* lines (Fig 1D & H). To further illustrate that NRC4’s perihaustorial accumulation pattern is not an optical artefact, we next co-expressed NRC4 C-terminally fused to orange fluorescent tag mKOk with GFP control and performed infection microscopy. In agreement with the results obtained when NRC4-GFP was expressed alone (Figure 1A), NRC4-mKOk produced a sharp fluorescence signal around the haustorium, unlike GFP control which displayed a uniform distribution pattern throughout the cell (Figure 1I & Movie S1). These results strongly suggest that NRC4 shifts its localization from cytosol to haustorium interface during infection.

**Figure 1.**
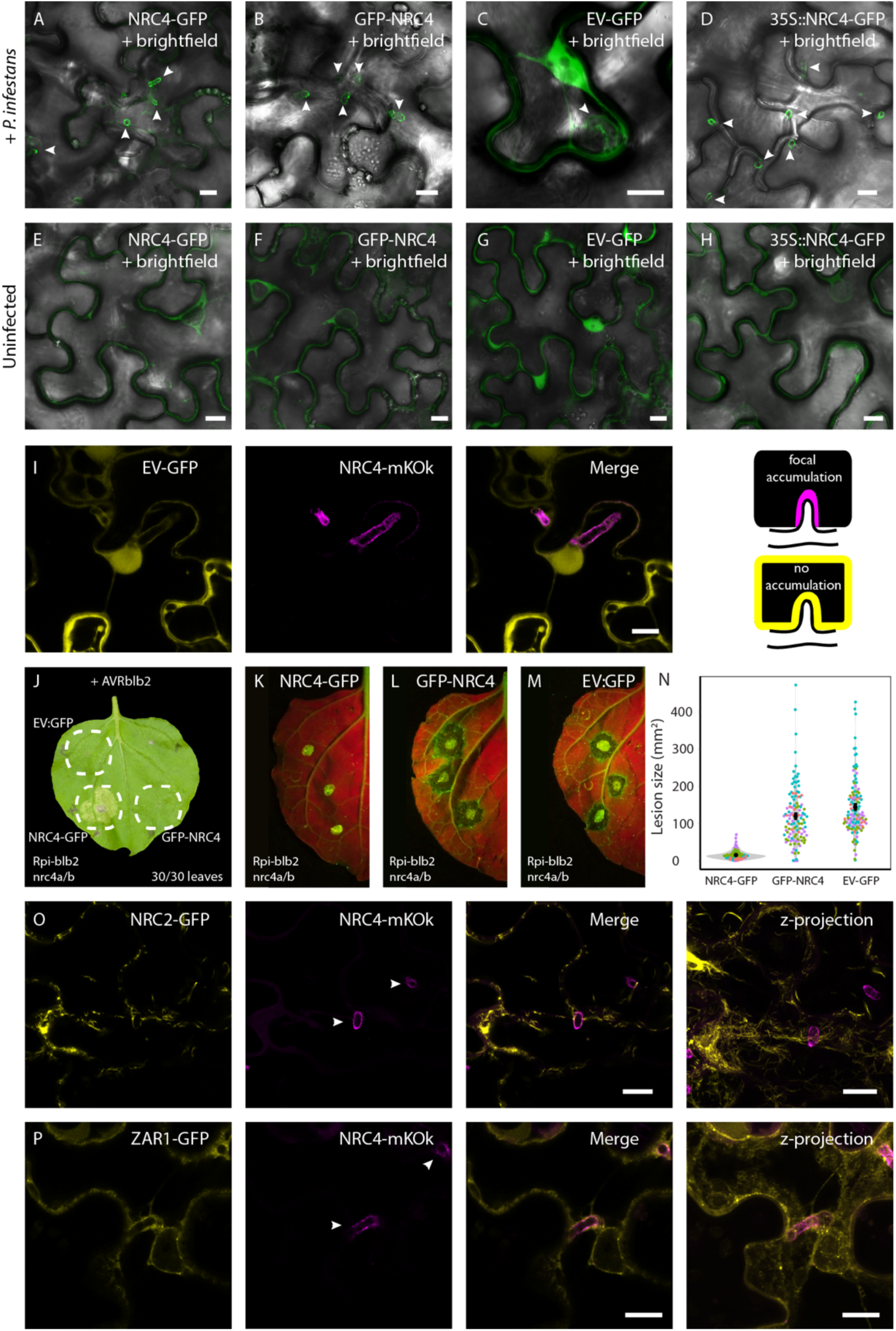
NRC4, but not NRC2 or ZAR1, accumulates at the EHM. (A-C) Single plane confocal micrographs showing Agrobacterium mediated expression of NRC4-GFP and GFP-NRC4 but not EV-GFP, accumulates around *P. infestans* haustorium. (D) Transgenic 35S::NRC4-GFP focally accumulates around pathogen haustoria. (E-G) Agrobacterium mediated expression of NRC4-GFP, GFP-NRC4 and EV-GFP all localise to the cytoplasm in uninfected cells. (H) Transgenic 35S::NRC4-GFP also cytoplasmic in uninfected cells. (I) Single plane images transiently expressing EV-GFP with NRC4-mKOk during infection with *P. infestans* showing EV-GFP does not focally accumulate around haustoria but localises to cytoplasm throughout the cell, including around haustoria, whereas NRC4-mKOk accumulates around haustoria. Cartoons (right) describe this result further. (J-N) NRC4-GFP but not GFP-NRC4 or EV-GFP genetically complement CRISPR/Cas9 mutation of NRC4a/b in Rpi-blb2 transgenic background by triggering (J) HR when co-expressed with AVRblb2 and (K-N) resistance when expressed alone and infected 1 day post agroinfiltration (dpai). (J) The result was replicated in 30 leaves, over three independent biological reps where *N* > 6 leaves per rep. Images were taken at 8 dpai. (K-N) Points in scatter violin plot (N) represent the area (mm^2^) occupied by infection spots measured on white light images, where each construct had *N* > 116 infection spots. The mean and standard error are shown as point and bars respectively. Four independent biological reps were conducted. UV (shown) and white light images were taken at 8 days post infection (dpi). Unpaired Wilcoxon tests gave p-values of 3.74×10^-33^ for NRC4-GFP to GFP-NRC4, 1.18 x 10^-37^ for NRC4-GFP to EV-GFP and 5.99 x 10^-3^ for GFP-NRC4 to EV-GFP. (O-P) Single plane images transiently expressing NRC2-GFP with NRC4-mKOk during infection with *P. infestans* showing NRC2 does not focally accumulate around haustoria but localises to disperse filaments. (Q) Z-projection of 10 z-slices. (R-T) Single plane images transiently expressing ZAR1-GFP with NRC4-mKOk during infection with *P. infestans* showing ZAR1 does not focally accumulate around haustoria, but is diffuse in the cytoplasm. (U) Z-projection of 11 z-slices. All scale bars are 10 μm.

In many cases, fluorescent fusions of NLRs lead to inactivity. Therefore, we next determined whether GFP fusions of NRC4 are functional. To do this we generated CRISPR/Cas9 mutants of NRC4a/b in the Rpi-blb2 transgenic background (hereafter Rpi-blb2^nrc4a/b^) and employed a HR complementation assay by co-expressing AVRblb2 with GFP fusions of NRC4 or an EV:GFP control in mutant plants. Infiltrated leaf patches on Rpi-blb2^nrc4a/b^ plants produced clear HR symptoms with C-terminally tagged NRC4 (NRC4-GFP) (n=30 plants, 100%) but not with EV:GFP (n=30 plants, 0%) or N-terminally tagged NRC4 (GFP-NRC4) (n=30 plants, 0%) (Fig 1J), showing that only the C terminal GFP fusion is functional. This was not due to reduced stability of GFP-NRC4 as this construct produced a stronger protein band compared to NRC4-GFP (Fig S1). We further validated that NRC4-GFP but not GFP-NRC4 or EV:GFP could genetically complement Rpi-blb2^nrc4a/b^ plants and provide resistance to *P. infestans*, (Fig 1K-N). These results are consistent with the finding that an N-terminal tag on ZAR1 inhibits its cell death function (32), possibly by blocking the α1 helix insertion into the membrane.

To determine if other NLRs can also accumulate around haustoria, or if this phenomenon is specific to NRC4, we investigated the localization of two other MADA motif containing NLRs NRC2 and ZAR1. We co-expressed NRC4-mKOk with GFP fusions of either the closely related helper NLR, NRC2, or the model CC-NLR, ZAR1 and performed infection microscopy. NRC2-GFP formed unusual filaments throughout the cell, some of which associated with the EHM, but did not focally accumulate (Fig 1N-Q and Fig S1). Likewise, ZAR1 did not accumulate around the haustorium but rather showed diffuse cytoplasmic distribution in haustoriated cells much like the GFP control (Fig 1R-U and Fig S1). These results demonstrate that not all NLRs accumulate at the haustorium interface like NRC4 and that NRC4 must have unique features or interactors governing haustorium targeting.

### NRC4 localizes to EHM microdomain(s)

The haustorium interface consists of several closely positioned compartments, namely: the plant cytoplasm, the EHM and the extrahaustorial matrix (EHMX) (12). To determine which haustorial interface compartment NRC4 localises to, we co-expressed NRC4 with various established marker proteins. We first co-expressed NRC4-mKOk together with GFP control in haustoriated cells. NRC4 fluorescence showed a sharp peak consistent with the EHM enveloping the pathogen haustorium (Fig 2A), unlike the GFP control which remained diffuse in the cytoplasm that surrounds the EHM, indicating that NRC4 accumulates at the EHM. The EHM consists of micro-domains and has a distinct protein and lipid composition from the plasma membrane (7, 8). We therefore co-expressed NRC4 with the plasma membrane marker protein RFP-Remorin1.3 (RFP-Rem1.3) which was reported to localize to EHM subdomains during infection by *P. infestans* (7). RFP-Rem1.3 partially but not fully co-localised with NRC4-GFP at the EHM, indicating NRC4 accumulates at EHM sub-domains. Meanwhile, we found that NRC2-GFP localised to dispersed filaments some of which contacted the EHM labelled by RFP-Rem1.3 (Fig S2A, Movie S2), and ZAR1-GFP showed exclusion from the EHM, instead localising to the cytoplasm surrounding the EHM (Fig S2B).

**Figure 2:**
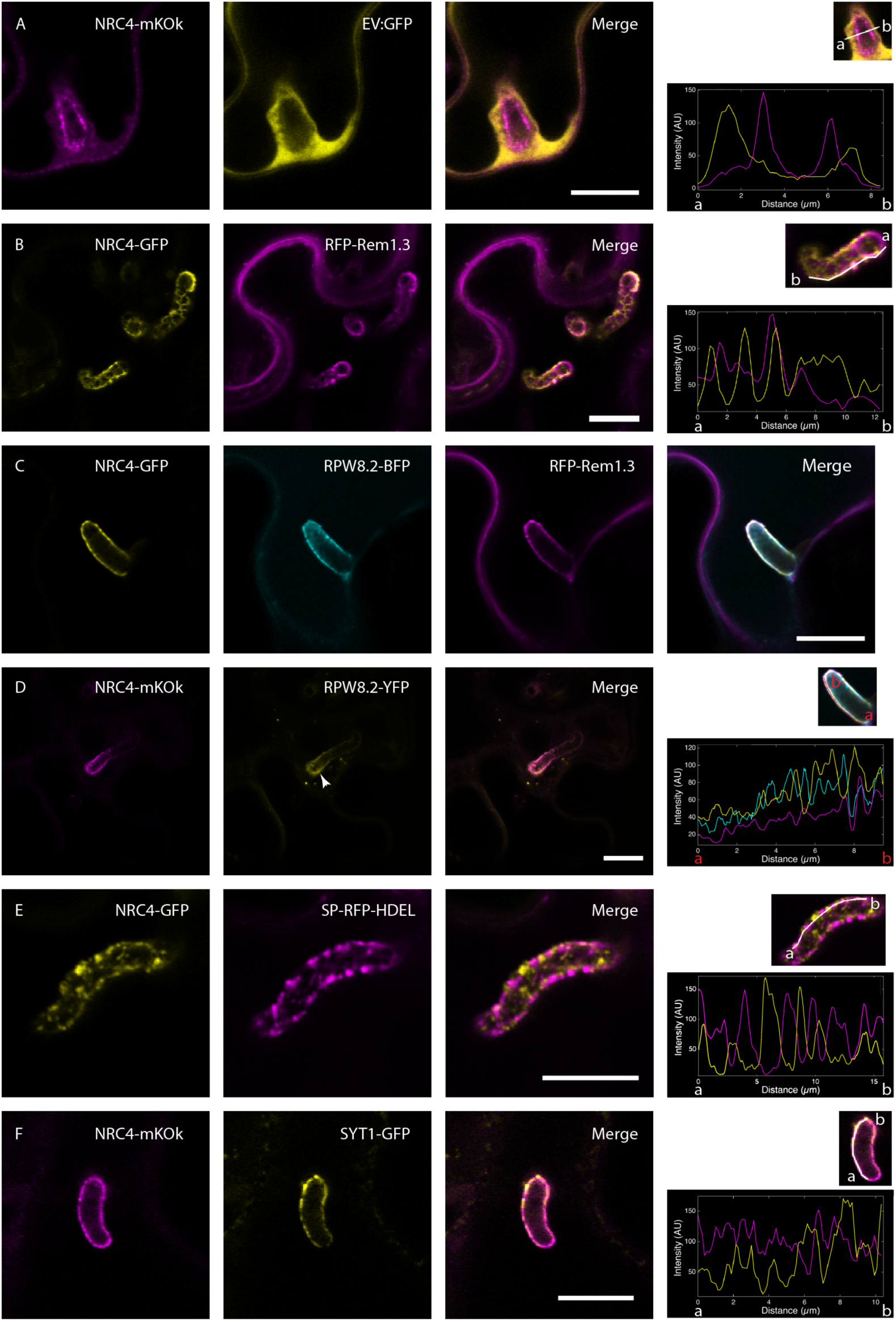
NRC4 localises to an EHM microdomain. (A) Peak NRC4-mKOk perihaustorial fluorescence does not overlap with the cytoplasmic marker EV-GFP. (B) NRC4-GFP shows a similar but not identical localisation to the PM and EHM marker *Solanum tuberosum* StRFP-Remorin1.3 (RFP-Rem1.3). (C) NRC4-mKOk shows a perihaustorial localisation partially overlapping that of EHM markers RPW8.2-BFP and RFP-Rem1.3. Line profile shown below merge image. (D) Some vesicles of RPW8.2-YFP are not labelled by NRC4, including one at the EHM (arrowhead). (E) NRC4-GFP perihaustorial signal is mutually exclusive to that of the ER marker SP-RFP-HDEL. (F) NRC4-mKOk shows a perihaustorial localisation mostly exclusive to that of GFP-SYT1. C was taken with Leica SP8, others SP5. Scale bars = 10 μm.

Because the Arabidopsis atypical resistance protein RESISTANCE TO POWDERY MILDEW8.2 (RPW8.2) was shown to accumulate at the EHM enveloping both powdery mildew fungi and the oomycete pathogen *Hyaloperonospora arabidopsidis* (33), we next checked its localization with regards to NRC4. RPW8.2-BFP showed a high degree of co-localisation with RFP-Rem1.3, and to a lower extent with NRC4 (Fig 2C) at the EHM during *P. infestans* infection of *N. benthamiana*. In addition, we also noted RPW8.2 mobile vesicle-like structures around the haustorium which are considered to be important for the transport of RPW8.2 to the EHM (34) (Movie S3). However, NRC4 did not localise to these RPW8.2-positive vesicles (Fig 2D arrowheads), suggesting that NRC4 might have a different transport route to accumulate at the EHM. Consistent with this, NRC4 is not predicted to have a secretion signal or transmembrane domain, unlike RPW8.2 which was reported to carry a predicted N-terminal transmembrane domain (35).

The endoplasmic reticulum (ER) surrounds the haustorium and associates with the EHM during fungal infections (11). Therefore, to rule out potential ER localization of NRC4, we co-expressed NRC4-GFP with the ER maker SP-RFP-HDEL. Although the SP-RFP-HDEL tightly wrapped around the *P. infestans* haustorium as reported during fungal infections, NRC4 did not co-localise with SP-RFP-HDEL (Fig S2C), excluding the possibility that NRC4 localizes to ER surrounding the EHM. Intriguingly, ER often appeared as puncta around the EHM (Fig 2E & S2D), resembling the swollen tubes of ER revealed around the fungal EHM (11). However, NRC4 was clearly excluded from these ER foci surrounding the EHM labelled by SP-RFP-HDEL (Fig 2E). In addition, we did not find a substantial overlap between fluorescence signals produced by NRC4-mKOk and Synaptotagmin1 (GFP-SYT1), a plasma membrane and ER associated protein which also accumulates at the EHM (Fig 2F). Collectively, these results show that NRC4 displays an uneven localization pattern across the EHM and is enriched in certain EHM microdomain(s).

### N-terminal α1 helix of NRC4 does not determine focal accumulation to the EHM

Accumulation of NRC4 but not closely related helper NLR NRC2 or the more distantly related MADA-containing NLR ZAR1 at the pathogen interface prompted us to study what part(s) of NRC4’s structure is required for EHM trafficking. To determine this, we first investigated the N-terminus of NRC4 as the α1 of ZAR1 that covers the MADA motif was reported to facilitate membrane association and act as a death-switch (22). We swapped NRC4’s α1 onto ZAR1 to see if ZAR1 could gain NRC4’s focal accumulation. However, ZAR1^NRC4α1^-GFP remained diffuse throughout the cytoplasm (Fig 3A) like WT ZAR1 (Fig 1R-U). Next, we performed the reciprocal swap, incorporating the α1 of ZAR1 to NRC4. Previously we named this chimera ZAR1_1-17_-NRC4 (23), hereafter named NRC4^ZAR1^*^α^*^1^-GFP. This chimera retained NRC4’s ability to accumulate at the EHM (Fig 3B), revealing that the first alpha helix does not determine NRC4’s EHM targeting. We confirmed our previous finding that this chimera was functional for HR and disease resistance (23), this time with Rpi-blb2^nrc4a/b^ plants (Fig 3C). Given the functionality of the chimeric NRC4, we hypothesised a shared mode of action of the α1 of ZAR1 and NRC4. We therefore built a model of NRC4 function based on ZAR1’s (Fig 3D-F). We suggest that NRC4 remains as a monomer or dimer in its resting state, adopting a closed inactive conformation in which the α1 helix remains unexposed as revealed by ZAR1 Cryo-EM structure. Following activation, NRC4 also probably oligomerizes and forms a resistosome, and the α1 helix flips out as in the case of activated ZAR1. Considering that the α1 helix is predicted to be buried in the coiled-coil domain (Fig 3D), this model is in agreement with our finding that the α1 helix does not determine the EHM localisation (Fig 3B). If this model is accurate, α1 would only be accessible for membrane association/protein interaction in the active state, but NRC4 accumulates to the EHM in the absence of a sensor NLR (Fig 1A), i.e. the inactive state.

**Figure 3.**
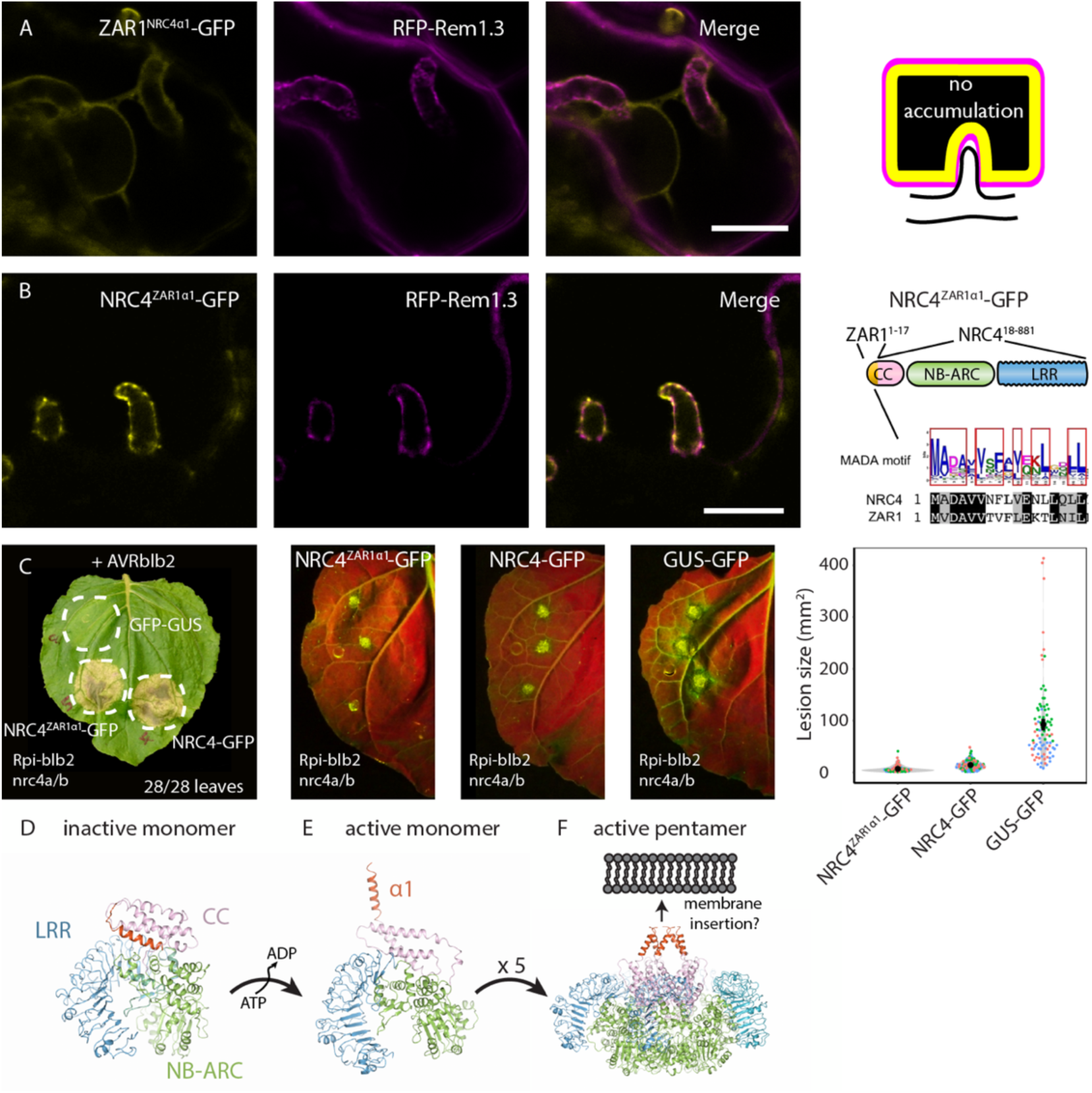
The first alpha helix of ZAR1 is functional in NRC4 and does not alter its localisation. (A) Single plane confocal micrograph showing ZAR1^NRC4α1^-GFP does not focally accumulate at EHM but is cytoplasmically localised. (B) Single plane confocal micrograph showing NRC4^ZAR1α1^-GFP focally accumulates at EHM with RFP-Rem1.3. Model depicts the chimeras of NRC4^ZAR1α1^. (C) NRC4 chimera with the N-terminal 17 amino acids from ZAR1, and NRC4-GFP but not EV-GFP genetically complements Rpi-blb2^nrc4a/b^ background by triggering HR when co-expressed with AVRblb2 and resistance when expressed alone and infected 1 dpai. The HR assay was repeated with the same results in 28 leaves, over two independent biological reps where N > 11 leaves. Images were taken at 8 dpai. Scattered points in scatter violin plot represent the area (mm^2^) occupied by infection spots measured on white light images, where each construct had N > 88 infection spots. The mean and standard error are shown as point and bars respectively. Three independent biological reps were conducted as indicated by colour of dots. UV (shown) and white light imaging was taken at 8 dpi. Unpaired Wilcoxon tests gave p-values of 1.33 x10^-10^ for NRC4-GFP to NRC4^ZAR1α1^-GFP, 1.84 x 10^-29^ for NRC4^ZAR1α1^-GFP to GUS-GFP and 6.99 x 10^-27^ for NRC4-GFP to GUS-GFP. (D-F) Homology model of NRC4 based on ZAR1 in the (D) inactive ADP-bound state (possibly monomeric), (E) newly active ATP-bound state (possibly monomeric) and (E) where five copies of NRC4 oligomerise into a pentameric resistosome and the five α1 helices insert into the membrane. CC domain (pink) consists of region 1-140 amino acids (aa), 141-157 is a disordered linker region, NB-ARC domain (green) 158-495 aa, LRR (blue) 496-843 aa.

### NRC4 requires a CC domain to accumulate at the EHM

We were able to use the homology model to determine precise domain boundaries and secondary structure boundaries of NRC4. We therefore investigated the intramolecular determinants of NRC4’s focal accumulation by truncating the N-terminus to remove the CC domain of NRC4 (NRC4ΔCC-GFP; NRC4Δ1-148-GFP). NRC4ΔCC-GFP lost its focal accumulation, but in some instances NRC4ΔCC-GFP formed puncta in the cytoplasm (Fig S3A). We quantified the enrichment of NRC4ΔCC-GFP at the EHM marker RFP-Rem1.3 versus the cytoplasmic marker EV-BFP, using image analysis (see methods). This revealed that NRC4ΔCC-GFP was not enriched at the EHM, unlike the full-length NRC4-GFP which produced a strong fluorescent signal spiking at the EHM (Fig 4). This indicated that NRC4 requires the CC domain for accumulation at the EHM. However, expression of NRC4’s CC domain alone (4CC hereafter; 1-148aa) fused to GFP did not show EHM accumulation (4CC-GFP), suggesting that the CC domain is necessary but not sufficient enough for haustorium targeting of NRC4 (Fig 4 & S3B). This finding also hinted that NRC4 carries additional features governing haustorium trafficking. We next swapped NRC4’s CC domain for NRC2’s and ZAR1’s. Both NRC4^2CC^-GFP and NRC4^Z1CC^-GFP accumulated at the EHM (Fig 4 & Fig S3C-D), suggesting NRC4 requires a coiled-coil domain but not necessarily its own.

**Figure 4:**
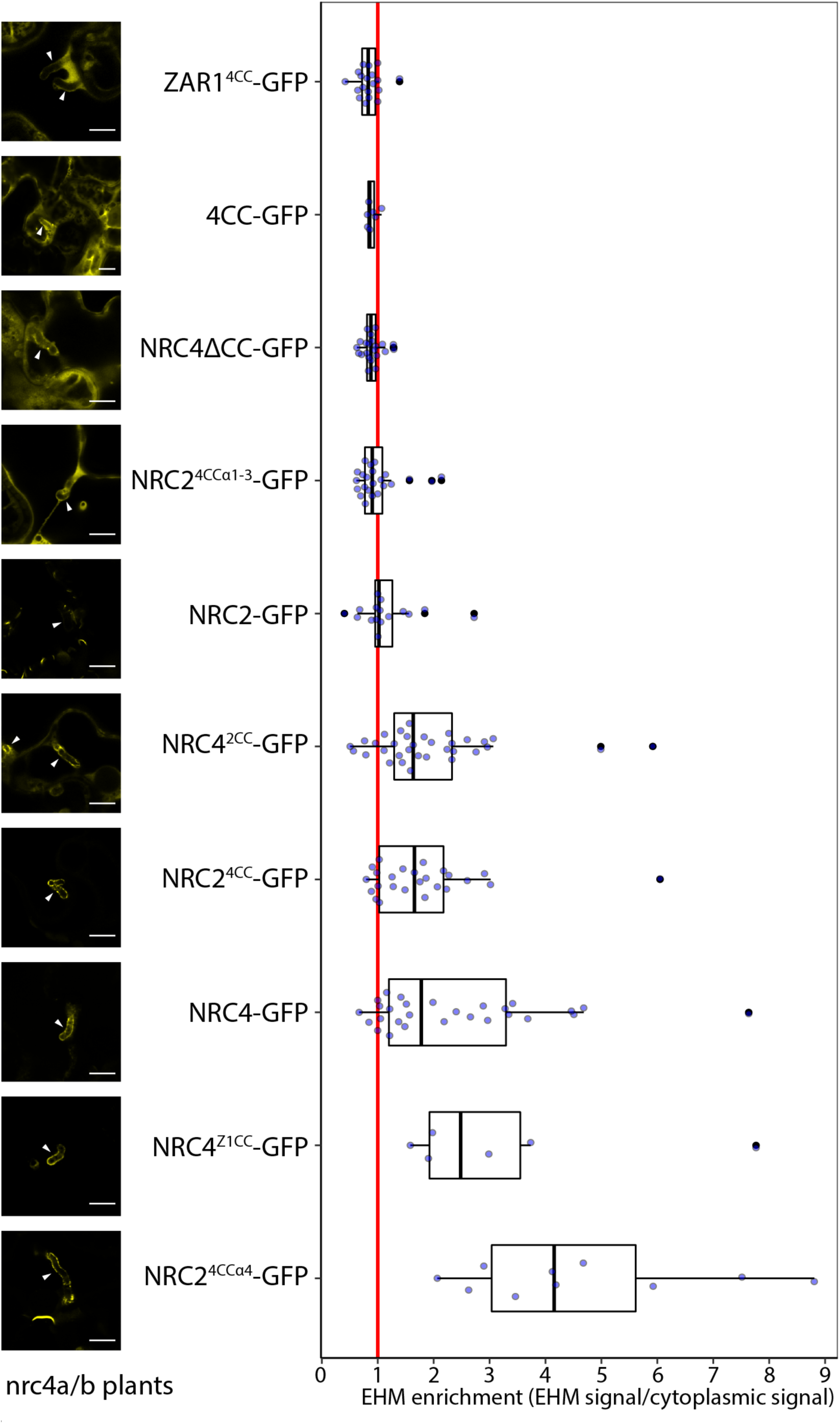
NRC4 requires a CC domain to accumulate to the EHM. NRC4, NRC2 and ZAR1 C-terminally fused GFP chimeras were co-expressed with plasma membrane and EHM marker RFP-Rem1.3 and cytoplasmic marker BFP-EV, and silencing suppressor P19 to boost expression. Leaves were infected and after three days were imaged with a Leica SP8 microscope. Single-plane micrographs were captured with the names of the constructs blinded to reduce acquisition bias, the image names were also randomised to reduce bias during quantification. A custom ImageJ/FIJI macro was made (see materials & methods) which allowed us to quantify the GFP signal at the peak RFP position (EHM) and divide it by the GFP signal at the peak BFP position. This is the EHM enrichment, and a number of 1 is no enrichment and a number of more than one is enrichment at the EHM. BFP and RFP channels are available in Fig S3.

Next, to see if NRC4’s CC domain could confer EHM targeting to ZAR1 or NRC2, we generated chimeras of these NLRs containing the CC domain of NRC4. A ZAR1-GFP construct carrying NRC4’s CC domain (ZAR1^4CC^-GFP) did not accumulate at the EHM (Fig 4 & S3E). In contrast, introduction of NRC4’s CC domain to the more closely related helper NLR NRC2 resulted in gain of function regarding EHM accumulation, as the NRC2^4CC^-GFP chimera displayed a clear EHM enrichment profile (Fig 4 & Fig S3F). This was an unexpected outcome given the results that NRC4’s CC domain alone cannot mediate haustorium enrichment when fused to GFP (CC-GFP) or introduced into ZAR1 (ZAR1^4CC^-GFP) backbone. However, these results further support our view that NRCs could carry additional structural features that mediate potential interactions with regulatory components for membrane trafficking.

We next investigated whether any specific region within NRC4’s CC domain (4CC) could mediate EHM accumulation when introduced into NRC2. Structural prediction revealed that NRC4’s CC domain encodes four a-helices. The first half of NRC4’s CC domain comprises α1-3 helices (amino acids 1-83), and the second half of 4CC (amino acids 84-148) comprises a4 and a long, disordered region. We first made intra-domain swaps of approximately half of 4CC onto NRC2 backbone. NRC2^4CCα4^-GFP gained NRC4’s focal accumulation at the EHM (Fig 4 & Fig S3G), whereas NRC4^2CCα1-3^-GFP did not (Fig 4 & Fig S3H). Although these findings implicate NRC4’s α4 helix in EHM enrichment, this data may seem counterintuitive given the EHM accumulation of NRC4 chimera carrying the CC domain of NRC2 (Fig 4 & S3C). However, since NRC2 is able to associate with the EHM to an extent (Fig 1O & Fig S3I), but does not accumulate (Fig 4), it is plausible that in addition to the CC domain NRCs foster multiple signatures that determine plasma membrane versus EHM localization. The balance between the activities of these different regions could determine subsequent membrane positioning of NRCs during infection.

### Activated NRC4 forms puncta that associate with both the EHM and plasma membrane

We next set to determine the fate of activated NRC4 in infected cells and compare it to the clear EHM accumulation of inactivated NRC4. To do this we needed to use an NRC4 mutant which can get activated but is unable to trigger cell death. Considering the first alpha helix of NRC4 is not involved in EHM targeting, we reasoned we could mutate this region to suppress HR without compromising NRC4’s ability to traffic to the EHM. The L9E mutant is predicted to interfere with α1’s ability to insert in the membrane and thus prevent HR (22, 23). Previously we showed that NRC4^L9E^ mutation suppresses HR and does not provide Rpi-blb2-mediated resistance during transient co-expression with Rpi-blb2 (23). We confirmed these observations in Rpi-blb2^nrc4a/b^ plants (Fig 5A), and also found that the L9E mutation did not compromise NRC4’s ability to traffic to the EHM (Fig 5B-C).

**Figure 5:**
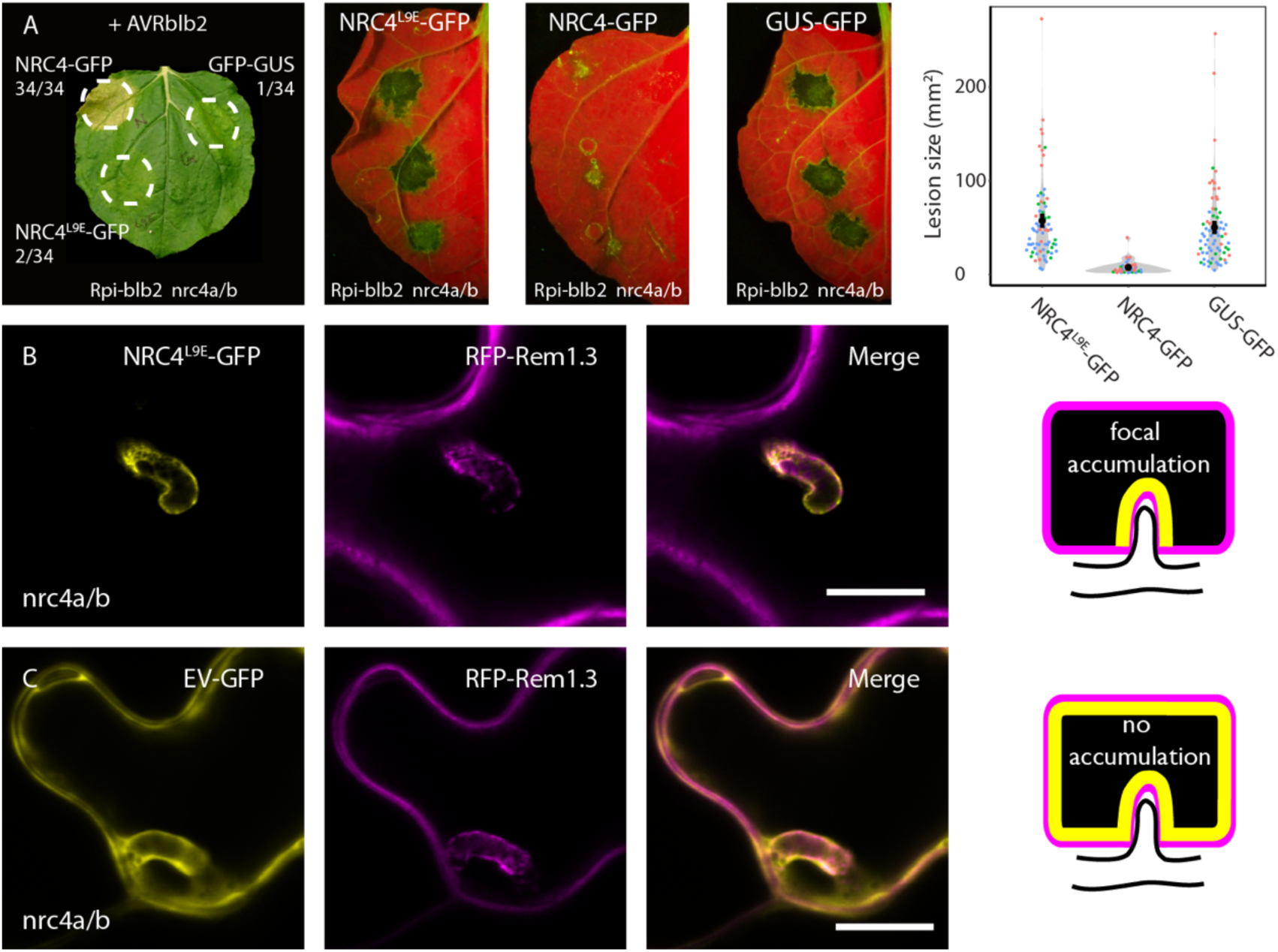
NRC4^L9E^ is non-functional for HR and *Phytophthora infestans* resistance but still focally accumulate at the EHM. (A) NRC4-GFP, but not NRC4^L9E^ mutant or EV-GFP genetically complements CRISPR/Cas mutation of Rpi-blb2^nrc4a/b^ plants by triggering HR when co-expressed with AVRblb2 and resistance when expressed alone and infected 1 dpai. The HR assay was repeated with the same results in 34 leaves, over two independent biological reps where *N* > 11. Images were taken at 3 dpai. Scattered points in scatter violin plot represent the area (mm^2^) occupied by infection spots measured on white light images, where each construct had *N* > 35 infection spots. Three independent biological reps were conducted as indicated by colour of dots. UV (representative images for three constructs) and white light imaging was taken at 8 dpi. Wilcoxon unpaired tests gave p-values of 5.00 x 10^-15^ for NRC4^L9E^-GFP to NRC4-GFP, 0.272 for NRC4^L9E^-GFP to GUS-GFP, and 1.46 x 10^-14^ for NRC4-GFP to GUS-GFP. (B-C) Single plane confocal micrographs showing NRC4^L9E^-GFP focally accumulates at EHM with RFP-Rem1.3, but not the EV-GFP control which remains cytoplasmic. Scale bars are all 10 μm.

Considering this mutant is functional in terms of its localisation (Fig 5B), we hypothesised that the L9E mutation renders the NLR defective in triggering HR but is otherwise functional. To simplify the study of NRC4’s active and inactive states we first investigated it in the absence of *P. infestans* infection. When in its inactive, resting state NRC4^L9E^-GFP localised to the cytoplasm in Rpi-blb2^nrc4a/b^ plants (Fig 6A). We next investigated the fate of NRC4^L9E^-GFP in its activated state, in the presence of an effector. To do this we co-expressed it with AVRblb2Δ8, a truncate of the AVRblb2 that has lost its virulence function, but can still trigger HR (14). In the presence of BFP-AVRblb2Δ8, activated NRC4^L9E^-GFP predominantly localised to puncta which frequently associated with the plasma membrane marked by RFP-Rem1.3 (Fig 6B).

**Figure 6:**
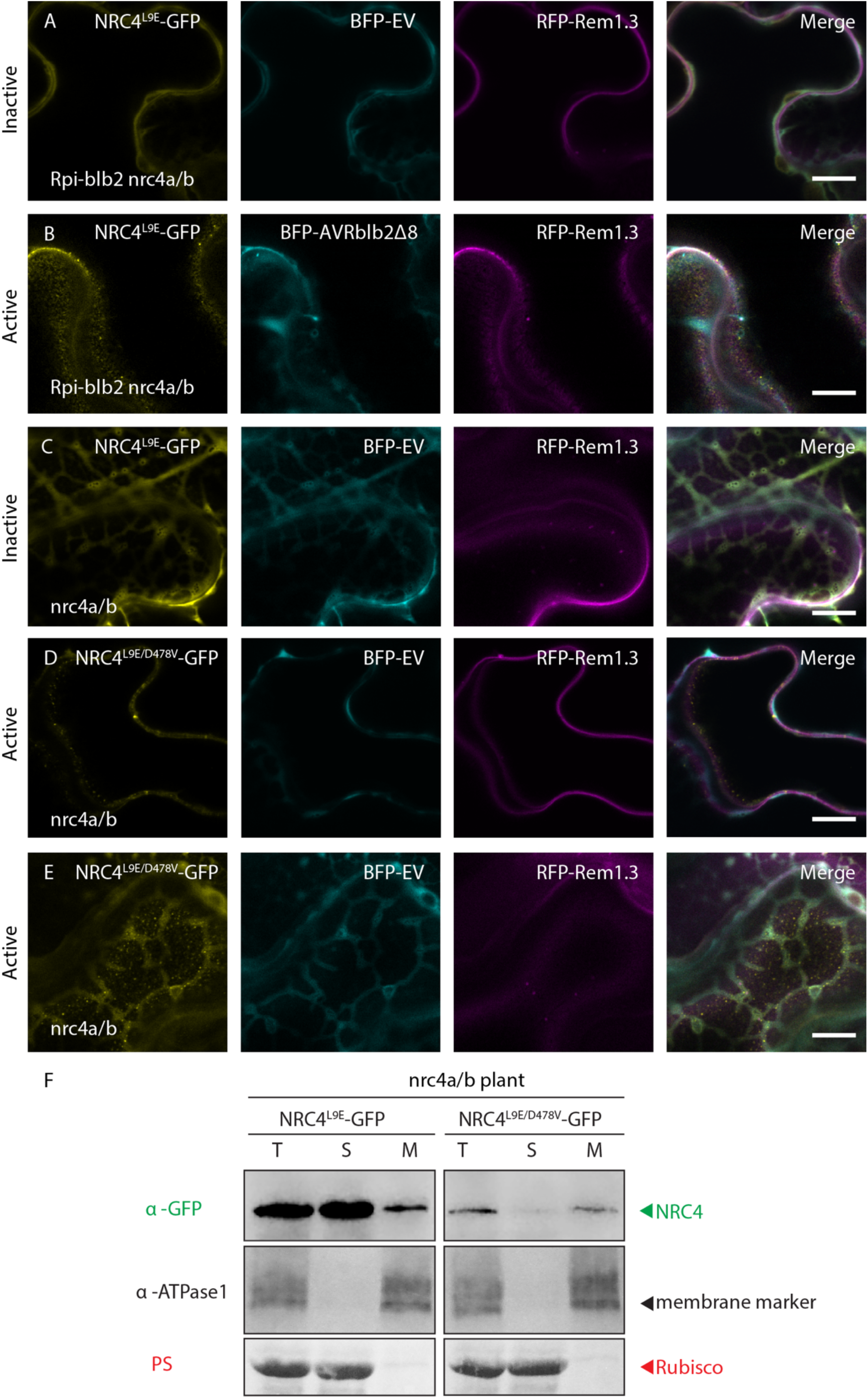
Activated NRC4 forms puncta associated with the plasma membrane in the absence of *Phytophthora infestans* infection. (A-E) Single-plane confocal micrographs showing the localisation of active and inactive variants of NRC4, with plasma membrane marker RFP-Rem1.3 and cytoplasmic BFP marker. All scale bars are 10 μm. (A) NRC4^L9E^-GFP is localized to the cytoplasm in Rpi-blb2^nrc4a/b^ plants when co-expressed with BFP-EV (B) NRC4^L9E^-GFP forms puncta associated with the plasma membrane when co-expressed with effector protein truncate BFP-AVRblb2Δ8. (C) NRC4^L9E^-GFP is localized to the cytoplasm when co-expressed with BFP-EV and RFP-Rem in nrc4a/b plants (D-E) Autoactive NRC4^L9E/D478V^-GFP forms puncta associated with the plasma membrane and patches of the cell labelled by both the cytoplasmic marker and membrane marker (F) Membrane enrichment confirms inactive NRC4^L9E^-GFP is mostly localized to the soluble (cytoplasmic) fraction, whereas autoactive NRC4^L9E/D478V^-GFP is more associated with the membrane fraction. T= total, S = soluble, M = membrane, PS = ponceau stain.

As an alternative to effector activation of NRC4, we used an autoimmune mutant of NRC4 by adding the D478V MHD mutation to the L9E HR suppressor mutant, *in cis* (NRC4^L9E/D478V^). In NRC4^L9E/D478V^, D478V renders NRC4 autoactive, but the L9E suppresses the cell death (23). We found that NRC4^L9E/D478V^-GFP localised to puncta in areas of the cell labelled by the membrane marker alone and areas of the cell where the cytoplasmic marker and membrane marker were present (Fig 6D-E). To validate these observations we performed membrane enrichment experiments in non-denaturing conditions, followed by SDS-PAGE (Fig 6F). We found that NRC4^L9E^ was present mostly in the soluble fraction, whereas NRC4^L9E/D478V^ was preferentially found in the membrane-enriched fraction. Lower protein levels of NRC4^L9E/D478V^ overall could be due to increased degradation of active NLRs as reported for RPM1 (31, 36). These biochemical findings that autoactive NRC4 associates more with membranes are in agreement with our microscopy that active NRC4 forms puncta which are mostly associated with the plasma membrane. It is possible that these punctate structures represent the oligomerized state of activated NRC4. However, it is unclear whether these are indeed NRC4 resistosomes, or perhaps groups of resistosomes enriched in membrane microdomains and further research is needed to clarify their nature.

Next, we investigated the subcellular dynamics of NRC4 when it is activated via the sensor NLR Rpi-blb2 upon recognition of AVRblb2, presumably secreted during *P. infestans* infection. To overcome HR cell death that could limit live cell imaging, we expressed NRC4^L9E^ mutant in Rpi-blb2^nrc4a/b^ plants or nrc4a/b control plants that do not express Rpi-blb2. Remarkably, NRC4^L9E^-GFP accumulated around haustoria both in Rpi-blb2 and control plants. However, in the presence of Rpi-blb2, NRC4^L9E^-GFP formed punctate structures across the EHM (Fig 7A-B & S4A). Additionally, in haustoriated Rpi-blb2^nrc4a/b^ plants, NRC4^L9E^-GFP often displayed clear labelling of the plasma membrane and puncta that is located at the plasma membrane (Fig 7A, S4A & Movie S4). In other cases, NRC4^L9E^-GFP did not accumulate at the EHM in Rpi-blb2^nrc4a/b^ plants, but instead there was equal labelling of the EHM and plasma membrane, and puncta localised at both the EHM and plasma membrane (Fig S4B). These results reveal that activated NRC4 does not exclusively accumulate at the haustorium, showing plasma membrane localization to an extent, and forms puncta that mainly remain associated with the EHM but also localise at the plasma membrane. Remarkably, autoactive NRC4^L9E/D478V^-GFP did not accumulate at the EHM at all, but instead produced fluorescent signal scattered throughout the infected cell similar to the EV-GFP control (Fig 7C-D). However, unlike GFP control, NRC4^L9E/D478V^-GFP mainly labelled the plasma membrane and frequently produced puncta that were mostly associated with the plasma membrane but also with the EHM to some degree (Fig 7C-D), indicating that activated NRC4 could redefine its membrane targeting route.

**Figure 7:**
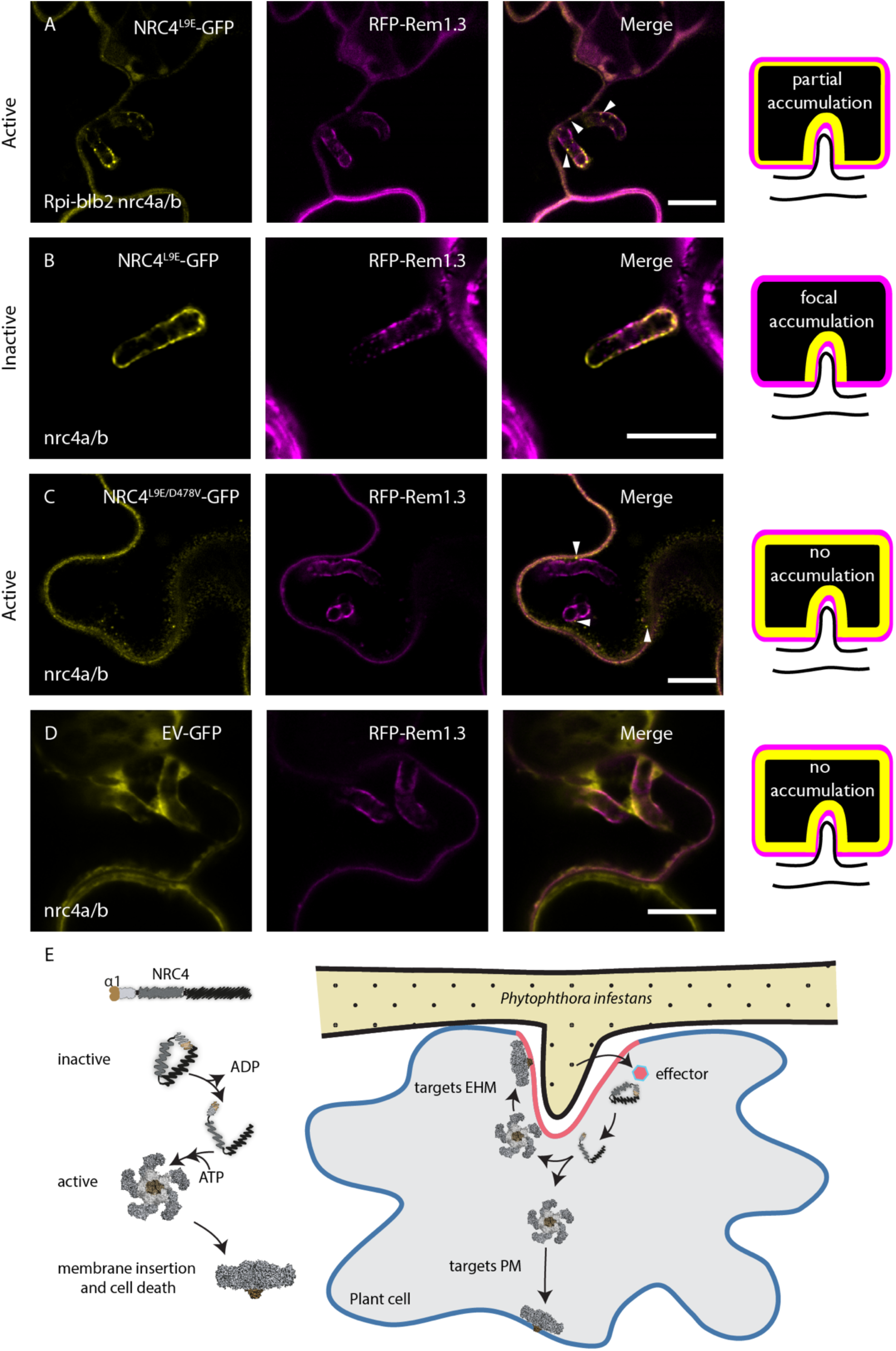
NRC4 is activated by Rpi-blb2 during infection and forms puncta that associate with the EHM and plasma membrane. (A-D) Single-plane confocal micrographs showing the localisation of active and inactive variants of NRC4, with plasma membrane marker RFP-Rem1.3 during infection with *P. infestans*. All scale bars are 10 μm. (A) NRC4^L9E^-GFP forms puncta associated with the EHM and plasma membrane in Rpi-blb2^nrc4a/b^ plants, and partially but not fully accumulates at the EHM. (B) NRC4^L9E^-GFP focally accumulates to the EHM in nrc4a/b plants. (C) NRC4^L9E/D478V^-GFP does not focally accumulate at the EHM, but instead forms puncta on the plasma membrane and EHM in nrc4a/b plants. (D) EV-GFP does not focally accumulate or form puncta in nrc4a/b plants. (E) Model depicting possible modes of action of NRC4 where NRC4 gets activated via sensor NLRs (not shown) at the EHM, by detecting perihaustorial effectors. NRC4 then oligomerises into a resistosome and targets the EHM, but a population of resistosomes dissociates from the extra haustorial membrane (EHM) to target the plasma membrane (PM). These resistosomes insert into the membrane to cause programmed cell death.

Based on these findings, we propose a model (EHM release model – Fig 7E) in which the non-activated NRC4 accumulates at the haustorium interface, positioning itself at effector delivery sites, possibly to improve its activation potential and to accelerate the deployment of the immune response. While activated NRC4 forms puncta associated with the EHM, some of the activated NRC4 is released from this initial site of activation to target the plasma membrane, possibly to propagate immune signalling and enhance the HR cell death response. Activated NRC4 could potentially form oligomers as illustrated by activated ZAR1 model, forming resistosomes that target the EHM and plasma membrane (Fig 7E).

## Discussion

Here, we identified an immune mechanism in which an NLR undergoes dynamic changes in its subcellular localization during infection, positioning itself at the specialized plant-pathogen interface where the corresponding pathogen effectors are deployed. Strikingly, following immune recognition, the activated NLR further re-organizes to form punctate structures not only at the pathogen interface membrane but also at the distant plasma membrane, suggesting that activated NLRs could possibly propagate from the primary site of activation to distant membrane interfaces. We report that NLRs are not necessarily stationary immune receptors, but instead can have inactive and active states shuttling to and from effector delivery sites.

Whether there is immune signalling at the EHM is currently unknown. In this work we uncovered that the inactive form of the helper NLR, NRC4 accumulates at the EHM surrounding the *P. infestans* haustorium. Considering the prominent roles of the EHM in facilitating effector translocation and nutrient uptake by the pathogen, as well as secretion of plant-defence compounds, it is not surprising that the pathogen deploys effectors to this site or that the plant deploys NLRs guarding it. Conceivably, positioning of receptors where their ligands accumulate would enhance their chances of recognition. Consistent with this view, two *P. infestans* effectors that accumulate at the haustorium interface, AVRblb2 and AVR1, are recognized by NRC4 dependent sensor NLRs Rpi-blb2 and R1 respectively (13, 14, 21). Intriguingly, both AVRblb2 and AVR1 are implicated in interfering with defense-related secretion (13, 14, 16). AVRblb2 interferes with secretion of an immune protease via an unknown mechanism, whereas AVR1 targets a key member of the secretory pathway, Sec5 of the exocyst complex at the EHM (13, 16). Therefore, we speculate that NRC4 could be guarding components of vesicle trafficking and fusion at the EHM by pairing with various sensor NLRs to monitor potential manipulation by effectors.

These findings implicate NLRs in plant focal immune responses. *P. infestans* is a useful model for studying focal immunity because of the large haustorial interface which allows clear identification of focal responses due to the substantial changes that occur upon haustorium penetration. However, focal immunity is not just restricted to haustoria forming pathogens (37–41). Therefore, our findings could also be relevant to non-haustoria forming pathogens and pests such as bacteria, nematodes, insects and viruses. Consistent with this view, NRC4 also pairs with sensor NLRs that recognize nematodes, insects, bacteria and viruses (21). Each has an interface with the host which can be targeted by plant focal immunity, and it would be interesting to determine if NLRs are repositioned in these pathosystems.

Here, we discovered that NRC4 but not closely related helper NLR NRC2 or the distantly related MADA NLR ZAR1 accumulates at the EHM (Fig 1O-P). Intriguingly, NRC2 formed fibril like structures some of which associate with the EHM, but overall it did not show accumulation like NRC4 (Fig 1O & S2A). Thus, to our knowledge, NRC4 represents the first case of a full length NLR that can target and accumulate at the specialized pathogen interface membrane, in plant or animal systems. Whether this could be applied to other NLRs remains to be determined. The atypical plant resistance protein RPW8.2 which lacks NBD or LRR domains was shown to localize to the EHM (33). However, how it contributes to immunity is still unknown. Some NLRs contain an RPW8-like N-terminal domain instead of a CC or TIR domain. These CC_R_’s include ADR1 and NRG1 which are now known to be helper NLRs for multiple TNLs and CC-NLRs (42–44). In addition, ADR1 and NRG1 are known to be required for resistance to several haustoria-forming pathogens (42, 44). Whether these helper NLRs also target the EHM would be interesting to determine. It could be that focal accumulation of NLRs to the EHM is a common resistance mechanism against haustoria forming oomycetes and fungi.

Considering our finding that NRC2 and ZAR1 do not focally accumulate at the EHM (Fig 1O-P), it draws into question how NRC4 achieves this feat. We found that NRC4 relies on a coiled-coil domain to enable focal accumulation at the haustorium interface, but that its coiled-coil domain alone is not enough for this process (Fig 4 & S3A-B). However, NRC2 gained focal accumulation from transfer of NRC4’s CC domain or even the second half of the CC domain (Fig 4 & S3F-G). This indicates that the α4 and/or the disordered region at the end of the CC domain contains one signature that determines focal accumulation. However, it’s likely another signature is required for focal accumulation and that this is found in NRCs but not all NLRs, as ZAR1 did not gain focal accumulation upon transfer of NRC4’s CC domain (Fig 4 & S3E). The requirement for an EHM targeting signature found in the nucleotide binding (NB) domain or a leucine rich repeat (LRR) domain fits our hypothesis that NRC4 does not traffic to the EHM via the same pathway as RPW8.2, bearing in mind that RPW8.2 doesn’t require a NB or LRR domain for EHM localisation. The EHM-targeting signatures of NRC4 may be structural or sequence specific and involve inter-molecular associations with trafficking pathways targeted to the EHM.

Intriguingly, we found that not only the pre-activated NRC4 but also the activated NRC4 shifts it localization pattern in response to infection. During infection of plants that carry the sensor NLR Rpi-blb2, we noticed perturbations in NRC4 accumulation at the EHM (Fig 7A & S4A-B), as we noticed additional NRC4-GFP signal at the plasma membrane. We recapitulated these findings using an autoactive mutant form of NRC4 which did not exclusively accumulate at the EHM, but instead mainly labelling the plasma membrane (Fig 7C). We also confirmed, with membrane enrichment experiments that NRC4 can increase its association with membranes upon activation (Fig 6F). These results revealed that activated NRC4 does not accumulate at the EHM, possibly because it could undergo conformational changes upon activation, leading to its dissociation from the EHM. Such a conformational change was reported for the MADA CC-NLR ZAR1, which assembles into a pentameric resistosome implicated in membrane targeting (22, 23). Whether ZAR1 mode of action could be applied to other MADA CC-NLRs still remains to be addressed. However, our previous and current work suggest that NRC4 mediated HR cell death and resistance can be maintained in NRC4 chimeras carrying the N-terminal α1 region of ZAR1 that was suggested to make membrane pores (Fig 3C) (23).

Strikingly, we discovered that activated NRC4-GFP forms puncta that associate with the PM in uninfected cells (Fig 6B & D-E). In agreement with our model of activation then EHM release, activation of NRC4 via a sensor NLR during infection also leads to formation of NRC4 puncta that associate with the EHM and plasma membrane (Fig 7A & S4A-B). These NRC4 puncta could emerge at and remain associated with the EHM following effector recognition, whereas some could be released from this primary site of activation to target the plasma membrane throughout the cell. Alternatively, activated NRC4 monomers could also disassociate from the EHM, get enriched at the PM and form puncta there, possibly to propagate immune responses throughout the cell and enhance the HR. Such repositioning of an NLR protein have also been reported following activation of the mammalian NLRP3 receptor. Prior to its activation, NLRP3 remains in the endoplasmic reticulum (ER) and cytosol, whereas activated NLRP3 assembles into inflammasome targeting various organelles, presumably to enhance perception of danger signals and enhance inflammasome assembly (45).

But what are these NRC4 puncta? One plausible explanation is that these are oligomers of NRC4 that assemble into resistosome like ZAR1. Whether the puncta presented here represent resistosomes or clusters of resistosomes remains to be determined. A recent preprint showed that autoactive NRG1.1 forms similar puncta when expressed in *N. benthamiana*. The authors correlated these puncta with pore formation and calcium influx, leading to cell death (46). These data would indicate that the active-NLR associated puncta are resistosomes, or at least groups of resistosomes. If validated, this method for identifying resistosomes by expressing HR-suppressed variants by mutation of their MADA-motif, coupled with live cell confocal microscopy, can be a quick and easy way to monitor resistosome formation, instead of challenging techniques such as Blue-Native PAGE, which is also not truly *in vivo*.

Here we introduced a cell biology dimension to study NLR function during live cell infection. So far, this has not been feasible due to cell suicide following NLR activation. We have established methods and tools to overcome this limitation (NRC mutant plants and point mutants that prevent HR but not trafficking) to employ live cell infection imaging of NLRs in haustoriated cells. Particularly, by investigating the trafficking of NRC4, a helper NLR that accumulates at the haustorium interface in a unique fashion, we provide novel insights into dynamics of NLR activation in infected cells. Dissecting the functional principles of plant NLRs and their cellular dynamics during infection are not only critical basic mechanistic studies but also will guide future studies to design synthetic NLRs that can boost immune activation at the pathogen interface.

## Supporting information

Dataset S1 Image Macro

## Acknowledgements

We thank Dr Sebastian Schornack, Dr Khaoula Belhaj and Dr Yasin Dagdas for the great comments and suggestions over the years. We thank Dr Joe Win for sending materials and for the discussions. We thank the funding agencies: Imperial Schrödinger PhD Scholarship (CD), Research England UK GCRF (CD), BBSRC DTP (EM, ZS, AYL), Gatsby Charitable Foundation (HA, CW, AM, SK). The Facility for Imaging by Light Microscopy (FILM) (SMR) at Imperial College London is part-supported by funding from the Wellcome Trust (grant 104931/Z/14/Z) and BBSRC (grant BB/L015129/1).

## Materials & Methods

### Growth and maintenance of plants

Wild type, mutant and transgenic *Nicotiana benthamiana* plants were grown in a controlled growth chamber at 25°C, in a mixture of 3:1 soil and vermiculite. Plants were kept under a high light intensity in long day conditions of 16 h light, 8 h night. Plants 3-4 week old plants were used for microscopy experiments, and 4-5 week old plants were used for infection assays.

### Phytophthora infestans growth, maintenance and infection of plants

*P. infestans* 88069 isolate was grown on rye sucrose agar media as previously described [Song 2009], in the dark at 18°C for 10-19 days before harvesting zoospores. Experiments assessing the susceptibility of leaf tissue under certain gene expression conditions, i.e. infection assays, were carried out on the abaxial (underside) of detached leaves as described previously (Song et al., 2009; Saunders et al., 2012). Infection of leaf material for the purpose of microscopy was carried out on the adaxial (topside) of leaves intact to the body of the plant, in humid conditions at 18°C. For all experiments, approximately 500-700 motile zoospores were used to inoculate the leaves, in cold water droplets of 10 μl per infection spots.

### Confocal microscopy

Leaf tissue was prepared for imaging by sectioning of desired area surrounding an infection spot using a cork borer size 4, and were mounted, live, in wells containing dH_2_O made in Carolina Observation Gel to enable diffusion of gasses. The underside of the leaf tissue was imaged, where stated, using a 63x water immersion objective lens on a Leica TCS SP5 resonant inverted confocal microscope or, where stated, a Leica SP8 with 40x water immersion objective. Laser excitations for fluorescent proteins on SP5 and SP8 are presented in Table S1 below. Z-stack and single plane images were later exported to Fiji (ImageJ) image analysis software (47) for image analysis and processing of contrast & brightness. Images shown are representative of more than one experiment and observations are from multiple cells and more than one leaf.

**Table S1:** Laser excitations on Leica SP5 and SP8 microscopes.

|  | SP8 (40x water objective) | SP5 (63x water objective) |
| --- | --- | --- |
| GFP | 488 nm (Argon) | 488 nm (Argon) |
| RFP | 561/594 nm (Diode) | 543 nm (He-Ne) |
| BFP | 405 nm (Diode) | 405 nm (Diode) |
| mKOk | 514 nm (Argon) | 543 nm (He-Ne) |

### Quantification of enrichment at EHM by image analysis

NRC4, NRC2 and ZAR1 C-terminally fused GFP chimeras were co-expressed with plasma membrane and EHM marker RFP-Rem1.3 and cytoplasmic marker BFP-EV, and silencing suppressor P19 to boost expression. Leaves were infected and after three days were imaged with a Leica SP8 microscope. Single-plane micrographs were captured with same laser (2%) and gain (100 V) settings for GFP constructs and same zoom factor (6.34), format 1024 x 1024, speed 200, line average 4. These settings were optimised to match the light microscopy resolution limit of ∼200 nm. We oversampled with a pixel size of ∼45 nm to visualise small objects clearly. The names of the constructs were blinded to reduce acquisition bias, the image names were also randomised to reduce bias during quantification (https://imagej.nih.gov/ij/macros/Filename_Randomizer.txt; Tiego Ferreira, 2009). A custom ImageJ/FIJI macro was made (attached) which allowed us to quantify the GFP signal at the peak RFP position (EHM) and the GFP signal at the peak BFP position. This was carried out by drawing a line (width 20) from the perihaustorial cytoplasm, across the EHM at a perpendicular angle to the EHM. These numbers were copied from ImageJ/FIJI to a spreadsheet and then the GFP signal at peak BFP position was divided by the GFP signal at peak RFP position to give the EHM enrichment. A number of 1 is no enrichment and a number of more than one is enrichment at the EHM.

### Transient gene expression and HR assays

Transient gene expression was achieved using agroinfiltration as described by Bos et al. (2006). *Agrobacterium tumefaciens* containing the desired plasmid was washed in water and resuspended in infiltration buffer (10 mM MES, 10 mM MgCl2 and pH5.6) and infiltrated into 4 week old Rpi-blb2^nrc4a/b^ *N. benthamiana* plants using needleless syringes. The optical density of the bacteria was OD_600_ = 0.3 for NRC4-related constructs and OD_600_ = 0.05 for 3xHA-AVRblb2. Images were taken 8 days post infiltration.

### Infection assays

Infection assays were carried out as described by (Song et al., 2009). Half leaves of NRC4a/b mutant in Rpi-blb2 transgenic *N. benthamiana* plants were infiltrated with OD_600_ = 0.3. Leaves were detached and infected 24h later and imaged 8 days post infection UV light. Quantification was carried out with the FIJI/ImageJ software (47) by measuring the area of the necrotic lesion of the pathogen. Statistical analysis was performed with R and used the ANOVA and Tukey test to assess significance. *UV imaging of infection assays:* Leaves were placed under direct illumination of two strong UV lamps as evenly as possible, in a dark room. A yellow coloured filter was attached to the camera lens and images were taken with long exposure of 2 seconds, F8.0 and ISO 400.

### Molecular cloning techniques

Molecular cloning into pK7WGF2 or pH7WGR2 derivatives was carried out as described previously (48). Briefly, plasmids were constructed with Gibson Assembly using the primers in Table S2 below, transformed into DH5α chemically competent *E. coli* cells by heat shock. Plasmids were amplified in *E. coli* and extracted by Promega Wizard Miniprep columns before being electroporated into *Agrobacterium tumefaciens* GV3101 electrocompetant cells. LB agar supplemented with Gentamycin and Spectinomycin was used to grow cultures carrying pk7WGF2/pH7WGR2. Sequencing was provided by Eurofins. We domesticated the Gibson Assembly vectors further by introducing a ccdB selection, which is excised by EcoRV during vector linearlisation.

To generate NRC4^L9E^-GFP, NRC4^L9E/DV^-GFP, ZAR11-17-NRC4-GFP (NRC4^ZAR1α1^-GFP), NRC2-GFP, ZAR1-GFP and GUS-GFP expression constructs, pCR8::NRC4^L9E^, pCR8::NRC4^L9E/DV^, pCR8::ZAR1_1-17_-NRC4 (Adachi et al., 2019), pCR8::NRC2 (Wu et al., 2016), pCR8::AtZAR1 (Harant et al., 2020, addgene no. 158487) and pICSL80016 [β-glucuronidase (GUS) with two introns, addgene no. 50332] without their stop codon were used as a level 0 modules for the Golden Gate assembly. The Level 0 plasmids were assembled with pICH85281 [mannopine synthase promoter+Ω (MasΩpro), addgene no. 50272], pICSL50034 (eGFP, TSL SynBio) and pICSL60008 [Arabidopsis heat shock protein terminator (HSPter), TSL SynBio] into the binary vector pICH47742 (addgene no. 48001).

**Table S2:**
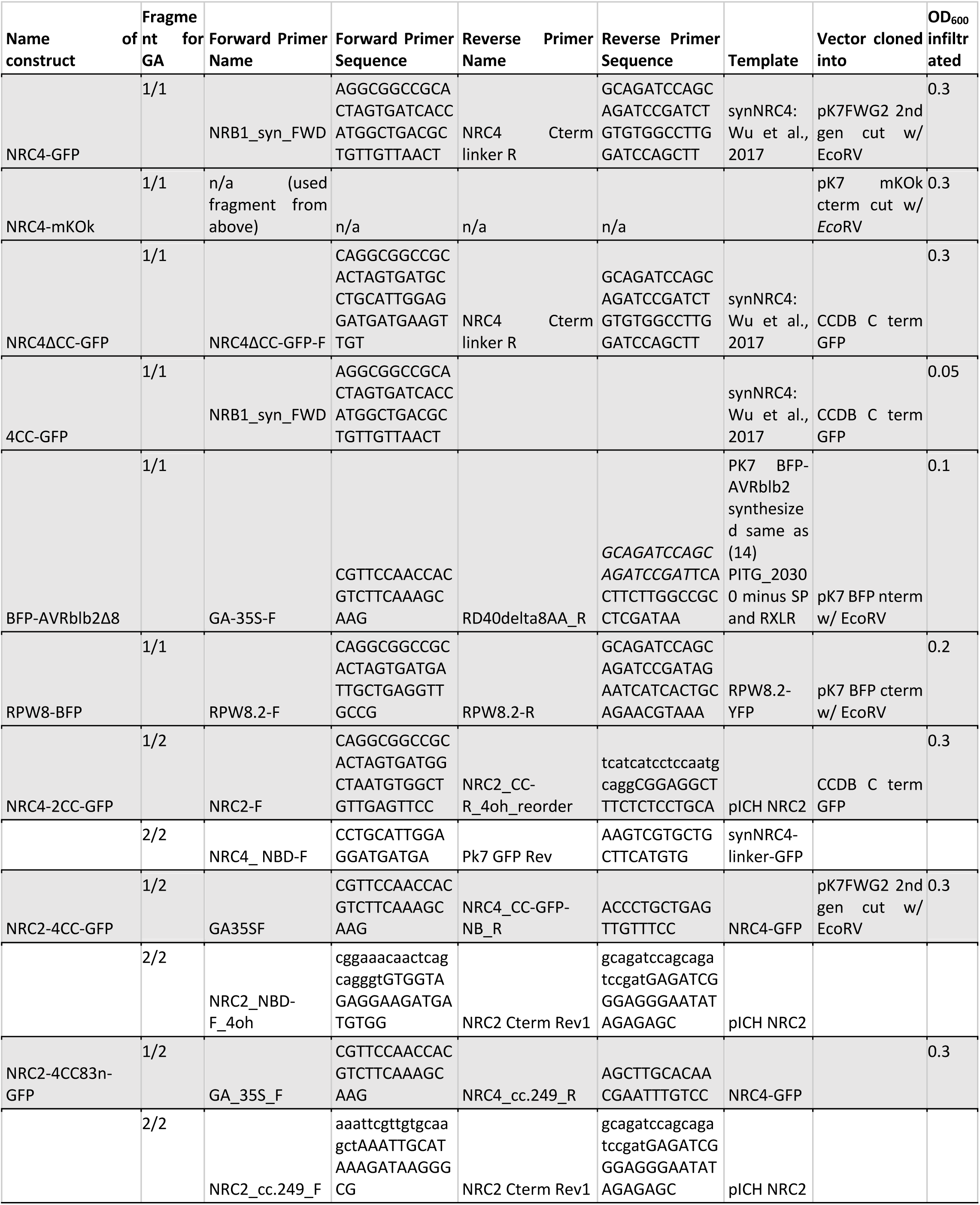
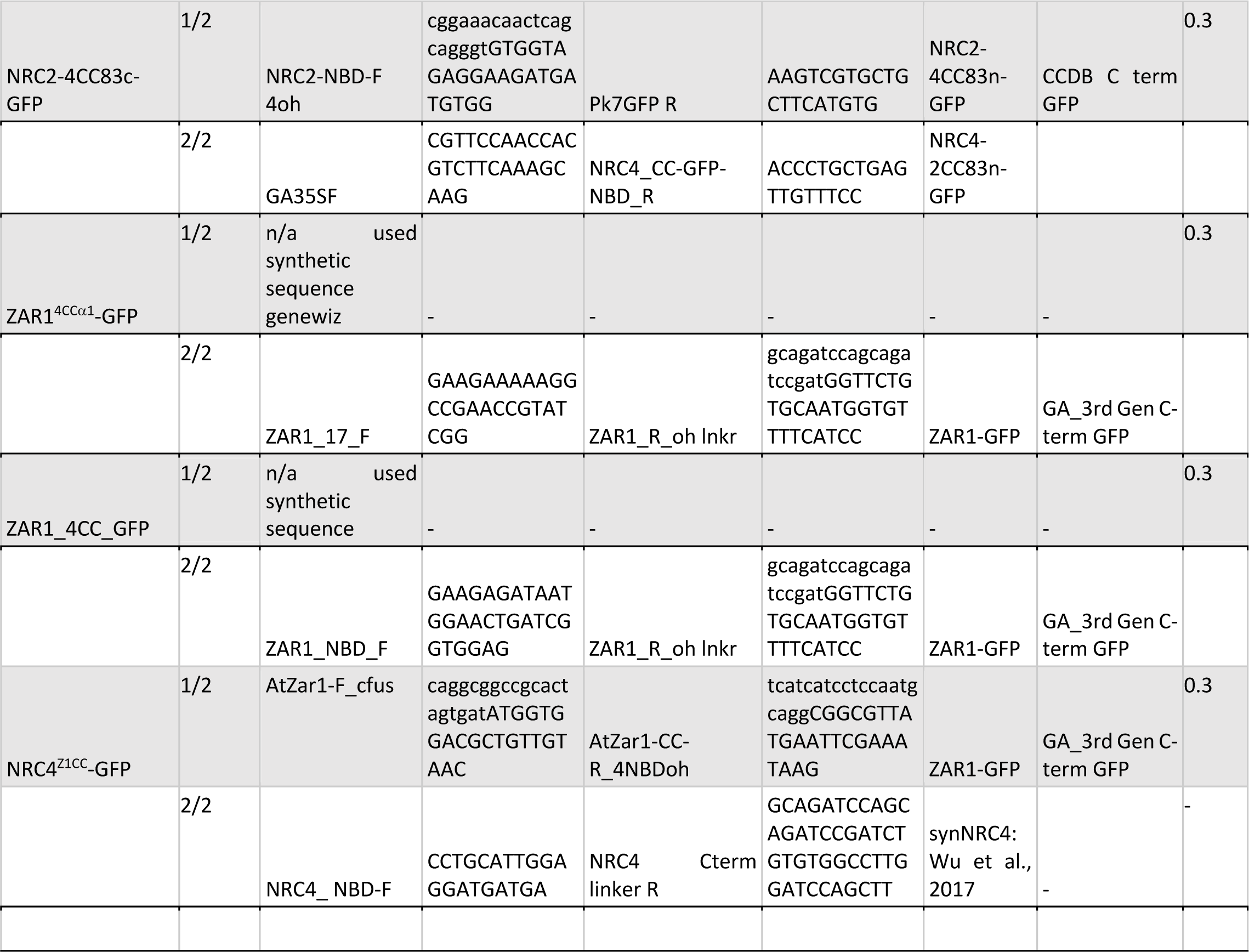
Details of new Gibson assembly constructs.

**Table S3:** Details of published constructs.

| Name of construct | Vector | Reference | OD<sub>600</sub> infiltrated |
| --- | --- | --- | --- |
| RFP-Rem1.3 | PK7WGR2 | (7) | 0.2 |
| GFP-SYT1 | PK7WGF2 | (7) | 0.2 |
| EV-GFP | PK7WGF2 | (Dagdas et al., 2016) | 0.1 |
| EV-3xHA | PK7 derivative | (Dagdas et al., 2018) | 0.1 |
| BFP-EV | PK7 derivative | (Dagdas et al., 2018) | 0.1 |
| RPW8.2-YFP | pPZPYFP | (33) | 0.2 |
| SP-RFP-HDEL | Vector unknown (Kan selection in bacteria) | (51) | 0.2 |

### Structure homology modelling

We used the cryo-EM structures of inactive and activated ZAR1 (25, 32) as template to generate homology models of inactive and activated NRC4. The amino acid sequence of NRC4 was submitted to Protein Homology Recognition Engine V2.0 (Phyre2) for modelling (49). The coordinates of inactive ZAR1 structure (6J5W) and activated ZAR1 resistosome (6J5T) were retrieved from the Protein Data Bank and assigned as modelling template by using Phyre2 Expert Mode. The resulting model of NRC4 comprised amino acids Val-5 to Glu-843 and was illustrated in CCP4MG software (50).

### Accession Numbers

The NRC4 sequences from this study are in the Solanaceae Genomics Network (SGN) or GenBank/EMBL databases with the following accession numbers: NbNRC4 (NbNRC4, MK692737; NbNRC4a, NbS00002971; NbNRC4b, NbS00016103).

### Membrane enrichment

Leaf tissue was collected and 600 mg per sample was frozen with liquid nitrogen then homogenised into a fine powder as with extraction for co-immunoprecipitation. Next, 2000 μl ice-cold sucrose extraction buffer was added (0.33 M sucrose, 20 mM Tris-HCL (pH 8), 1 mM EDTA) supplemented with 5 mM DTT, 1% Sigma Plant Protease Inhibitor and 0.5% PVPP. Samples were vortexed until they were liquid and well mixed. The lysate was cleared by centrifugation at 2000 x *g* for 5 min at 4°C to remove debris. Supernatant was filtered with two layers of Miracloth (carefully ensuring not too much of the extract was lost to the Miracloth due to absorption). An aliquot of this (50 μl) of this was separated into a new tube as the total fraction (T). The rest was spun at 21,000 x *g* (unless otherwise stated) for 1 h at 4°C, this was sufficient to enrich the membranes as despite popular belief, ultracentrifugation is not needed to enrich membranes if small sample volumes are used and the initial clearance is carried out at lower speeds than standard as to not pellet the heavy domains of membranes (52). The resulting supernatant was separated into a new tube and labelled as the soluble fraction (S). The pellet was resuspended with extraction buffer (minus the PVPP) in 3 times less volume than the soluble fraction (unless otherwise stated). The resulting fraction was labelled as the 3x membrane-enriched/microsomal fraction (M). Proteins fractions were run on SDS-PAGE gels as with western blotting, but with no water added to the loading buffer.

### Crude protein extraction and western blotting

Protein extraction, SDS-PAGE and western blotting was conducted as previously described (Dagdas et al., 2016). Whole or half-leaves were *Agro-*Infiltrated. Plants were incubated for 3 days at standard growth conditions to allow protein expression. Leaves were detached and 6 leaf disks were cut using cork borer tool size 4, and snap-frozen in 1.5 ml Eppendorf tubes in liquid nitrogen. Disks were stored at −80°C until protein extraction. *SDS-PAGE gel preparation:* Bio-Rad gel moulds were assembled and 1 mL autoclaved dH_2_O added to determine if there were any leaks. Water was removed by carefully tipping and air drying. The separating gel of 7.5% BIS-Acrylamide was prepared.

The separating gel was carefully pipetted into the mould, leaving 2-2.5cm space for the stacking gel. Isopropanol (0.5 ml) was added to remove any bubbles and the gel left to polymerise for an hour and before removing the isopropanol by carefully pouring and air drying. For gradient gels, 2.2 ml of 4% gel was aspirated into a 5 ml serological pipette followed by 2.2 ml of 16% gel. An air bubble was drawn up through the two gels and the mixture quickly pipetted into the glass mould. The stacking gel was then prepared using the following recipe for two gels: 1.25 ml 0.5 M Tris pH 6.8, 0.375 ml 40% acrylamide, 3.3 ml glass distilled water, 0.05 ml 10% SDS, 0.05 ml 10% APS, 0.005 ml TEMED. The stacking gel was pipetted up to the top of the glass plates and the combs were added, taking care not to allow any bubbles to be trapped in the wells/under the combs. This was allowed to solidify, and after the proteins were extracted the gel was assembled into the gel tank with 1x SDS running buffer.

#### Protein extraction

Tubes were kept in liquid nitrogen and not allowed to defrost throughout the process, until the step where extraction buffer was added. Leaf disks were ground to a fine powder evenly with a grinding tool and 120 μl extraction buffer added (30 μl Laemmeli sample buffer 4x [Bio-Rad], 24 μl DTT 1 M, 66 μl dH_2_O). Tubes were vigorously vortexed for 90 seconds, ensuring the buffer had reached all the tissue.

Tubes were centrifuged for the minimum amount of time possible in a microcentrifuge to bring tissue down from the sides, followed by denaturation of extracts at 80°C in preparation for SDS-PAGE. The tubes were centrifuged at 16,000 *g* for 5-10 minutes to pellet the tissue and allow the crude protein extract to remain in the supernatant. This supernatant was carefully removed to new tubes, avoiding leaf material and frozen at −20°C or left on ice until gel loading. The volume of protein extract to load depended on the experiment – unless otherwise stated in the figure legend, 19 μl was loaded from a 20μl master mix in order to account for pipetting error and evaporation. PageRuler Prestained Protein Ladder (4 μl) [Thermo Fisher] was added to one lane. Protein samples are carefully loaded and run in Bio-Rad Mini-PROTEAN Tetra Vertical Electrophoresis Cell using Bio-Rad PowerPac basic at 150 V constant until the proteins entered the separating gel, at which point the gel was run at 180 V until the ladder was separated enough to resolve the proteins of interest. *Western blotting:* PVDF membranes were activated in methanol for 60 seconds before equilibrating in 1X semi-dry transfer buffer [Bio-Rad] with the gel and transfer stacks (two stacks of 6-ply Wypall X60). The Trans-Blot Turbo Transfer System [Bio-Rad] was assembled (transfer stack -> membrane -> gel -> transfer stack) and the transfer machine set at 1.3 A, 25 V for 13 minutes to ensure transfer of low expressed, high-molecular weight proteins. *Detection:* Membranes were blocked for 1 hour shaking in 3% BSA blocking solution (in TBS), to prevent the primary antibody binding non-specifically. Appropriate primary antibody was added to in antibody solution (3% BSA solution in TBS supplemented with 0.1% Tween) and incubated shaking at 4°C overnight. Membranes were then washed with 1X TBS supplemented with 1% Tween 5 times for 5 minutes per wash, before adding the appropriate secondary antibody for 90 minutes. Blots were then washed as before. Luminol was then added for detection by chemiluminescence [ThermoFisher], or 5-bromo-4-chloro-3-indolyl phosphate (BCIP) and nitroblue tetrazolium (NBT) for colourimetric detection [Bio-Rad], depending on the conjugation to the particular secondary antibody, Horseradish Peroxidase (HRP) or Alkaline Phosphatase (AP) respectively. Chemiluminescence was recorded by LI-COR Odyseey Fx.

### Generation of transgenic and gene edited plants

Generation of Rpi-blb2^nrc4a/b^ CRISPR/Cas9 mutated plants was carried out the same as (23), where the same guide RNAs were used to target NRC4a/b. A T2 line unresponsive for HR upon transient expression with AVRblb2, but capable of complementing HR upon co-expression with NRC4-GFP was used for all experiments.

Generation of *35S::NRC4-GFP* plants was carried out as follows: Five week old WT *N. benthamiana* were infiltrated with *Agrobacterium* expressing NRC4-GFP supplemented with acetosyringone. At 3 days post agroinfiltration (dpai) leaves were collected and cut into 1-2 cm^2^ squares with a sterile scalpel. The largest veins were discarded. Leaf patches were surface sterilised in 70% EtOH for 30 s followed by 1.5% sodium hypochlorite solution with 5 drops of Tween20 surfactant per 500 mL solution. After this, the leaf patches were washed in sterile water 4 times. Each leaf square was placed adaxial side up in selective media (Table S4 below) and gently pressed to increase surface contact with the media. These plates were sealed with micropore tape and incubated in a plant transgenesis room at 22°C. Every two weeks patches were transferred to fresh plates. After 1 to 2 months shoots formed and were transferred to rooting media. After 4 weeks roots had formed and plantlets were transferred to soil in a high humidity chamber. Each day a little humidity was released until the T_0_ plantlets had adjusted to the standard growth chamber conditions. Plants were screened using confocal microscopy after a few weeks and those with GFP fluorescence were kept for seed. T_1_ seed was collected separately from their T_0_ parent. T_1_ lines with the best fluorescence were kept. T_2_ seed was collected from individual parent plants separately. A T_2_ line was selected for experiments as being the best expression (tNRC4-GFP #1.3).

**Table S4:**
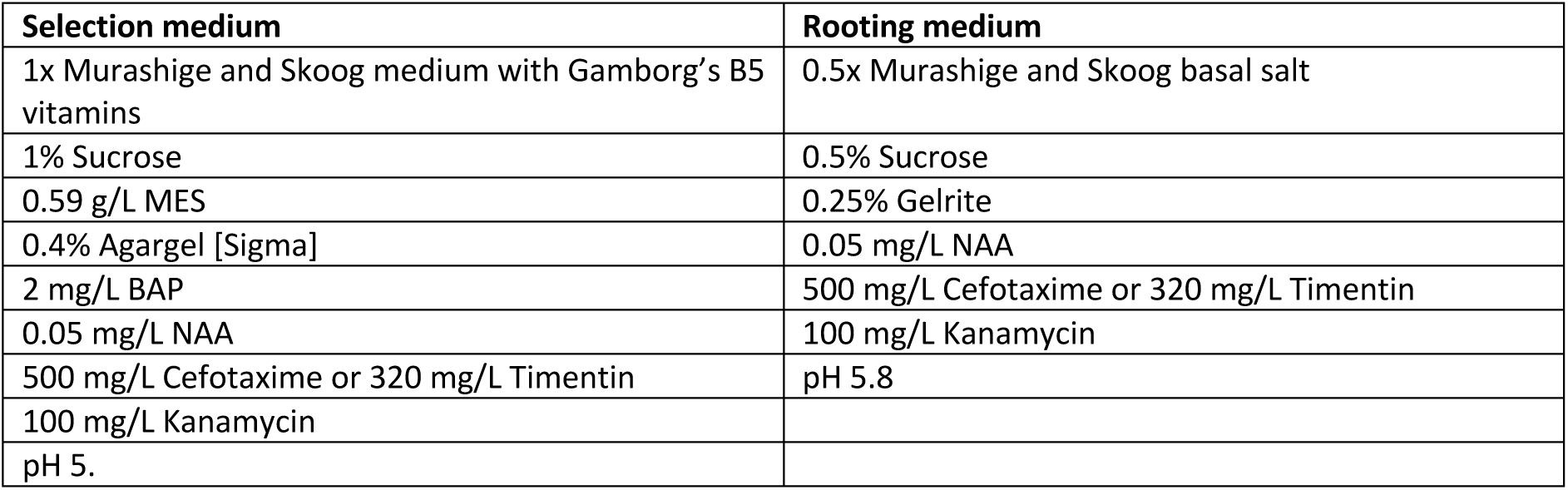
Composition of shooting and rooting media.

| **Selection medium** | **Rooting medium** |
| --- | --- |
| 1x Murashige and Skoog medium with Gamborg's B5 vitamins | 0.5x Murashige and Skoog basal salt |
| 1% Sucrose | 0.5% Sucrose |
| 0.59 g/L MES | 0.25% Gelrite |
| 0.4% Agargel [Sigma] | 0.05 mg/L NAA |
| 2 mg/L BAP | 500 mg/L Cefotaxime or 320 mg/L Timentin |
| 0.05 mg/L NAA | 100 mg/L Kanamycin |
| 500 mg/L Cefotaxime or 320 mg/L Timentin | pH 5.8 |
| 100 mg/L Kanamycin |  |
| pH 5. |  |

## Supplementary Figures

**Fig. S1.**
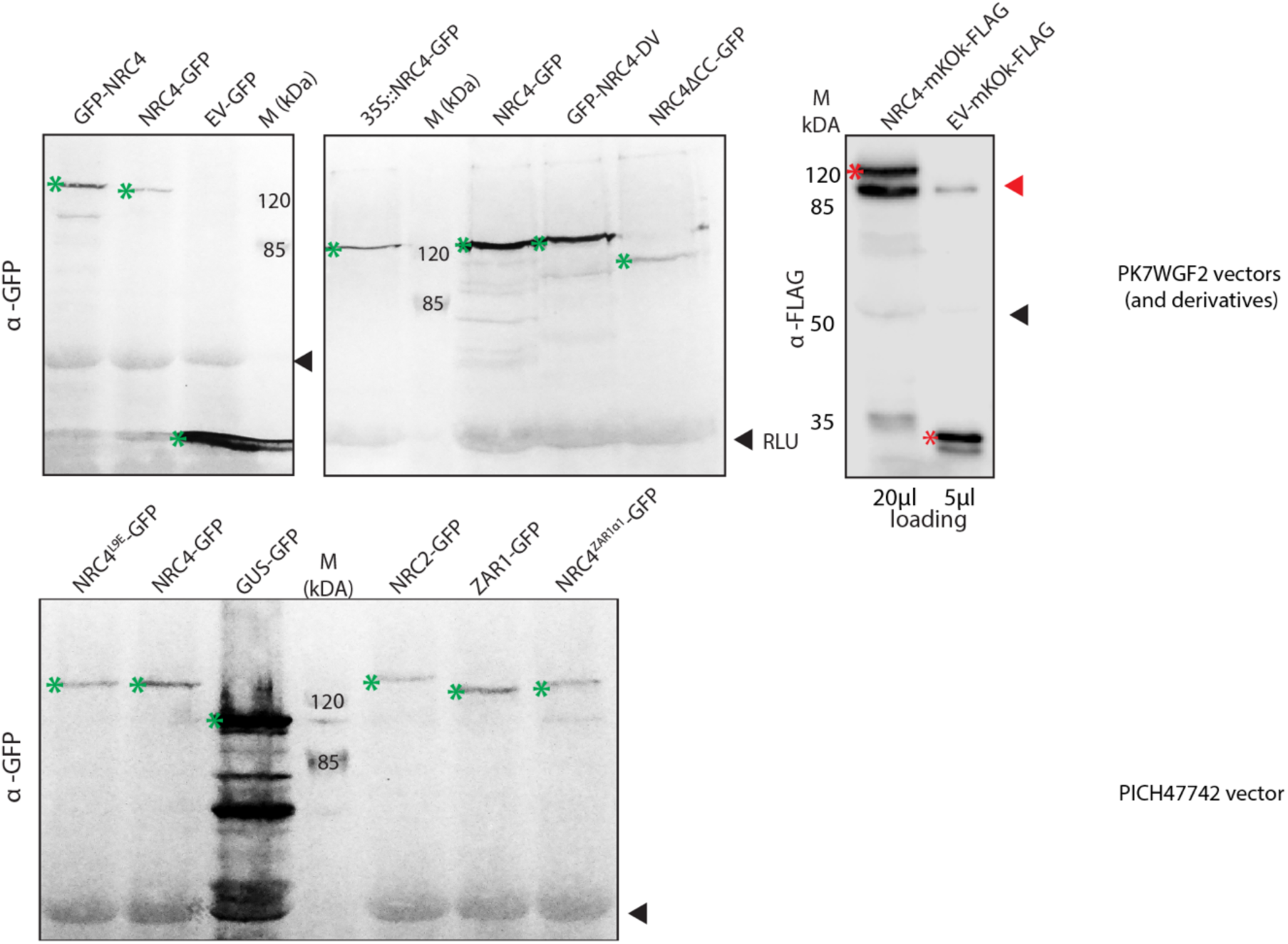
Fusion proteins of NLRs are expressed. Western blots showing expression of GFP or mKOk-FLAG tagged fusion proteins at 3 days post agroinfiltration (dpai). Top blots are of proteins expressed from pK7WGF2 vectors or its derivatives, bottom blot is of pICH47742 vector. RLU is rubisco large subunit stained by alkaline phosphatase-mediated colour detection of proteins. Asterisks indicated expected bands. Black arrowheads indicate rubisco large subunit, red arrowhead indicates non-specific band from anti-FLAG.

**Fig. S2.**
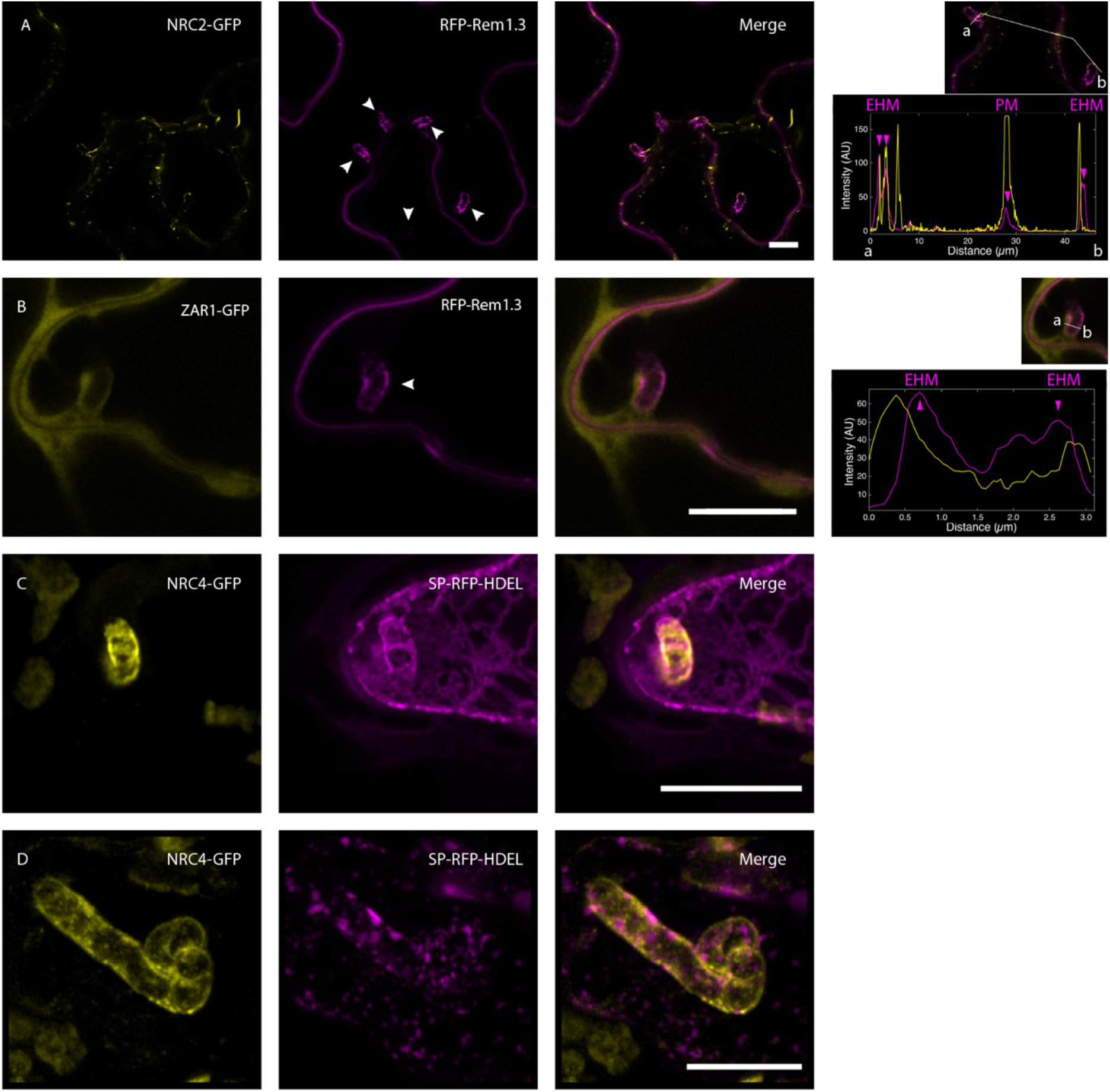
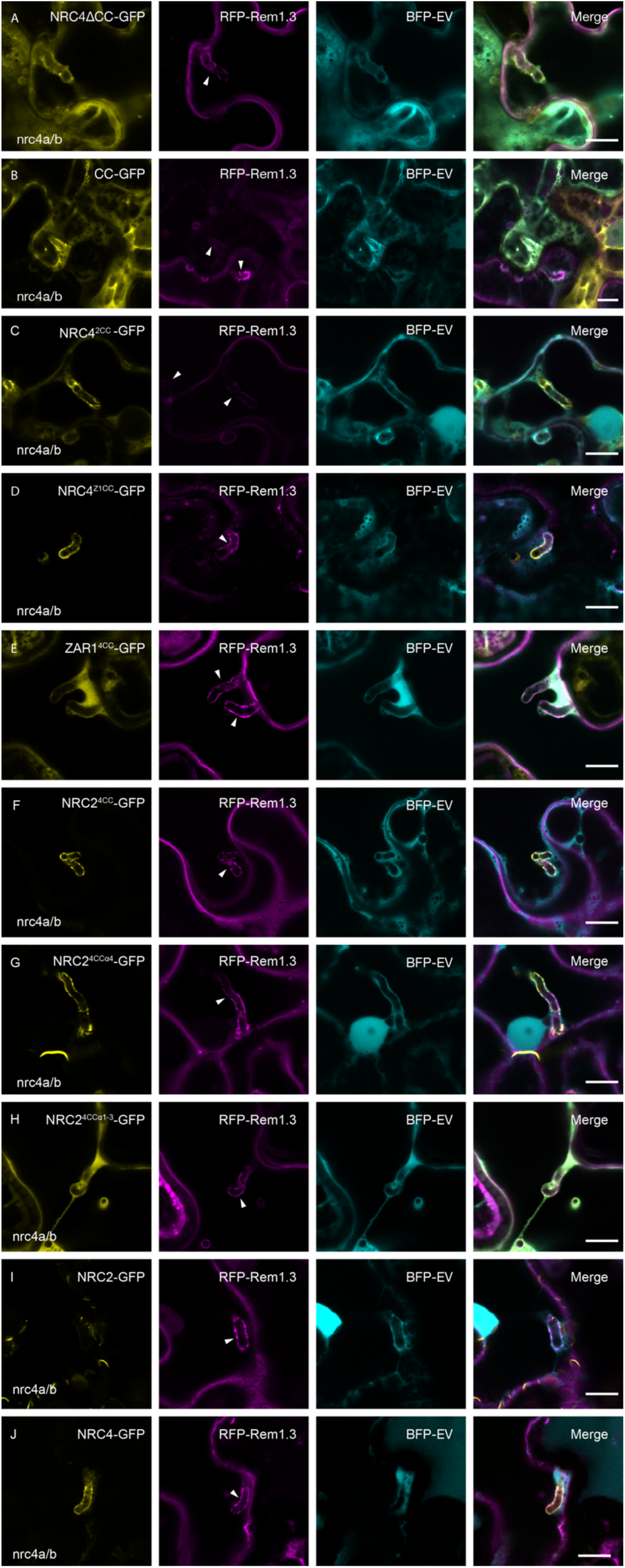
NRC2 and ZAR1 do not accumulate at the EHM with RFP-Rem1.3 and NRC4 does not substantially co-localise with the ER marker SP-RFP-HDEL. (A-B) Single plane confocal micrographs. (A) NRC2-GFP forms filaments throughout the cell, some of which associate with or overlap with the plasma membrane (PM) and extrahaustorial membrane (EHM) marker; but NRC2 does not focally accumulate at the EHM. (B) ZAR1-GFP does not focally accumulate at the EHM with RFP-Rem1.3, but instead is diffuse throughout the cytoplasm (C-D) Z-projections of 45 and 61 z-slices respectively. NRC4-GFP perihaustorial signal is mutually exclusive to that of the ER marker SP-RFP-HDEL. A-B were taken with Leica SP8, C-D were taken with Zeiss Airyscan. Scale bars = 10 μm.

**Fig. S3.**
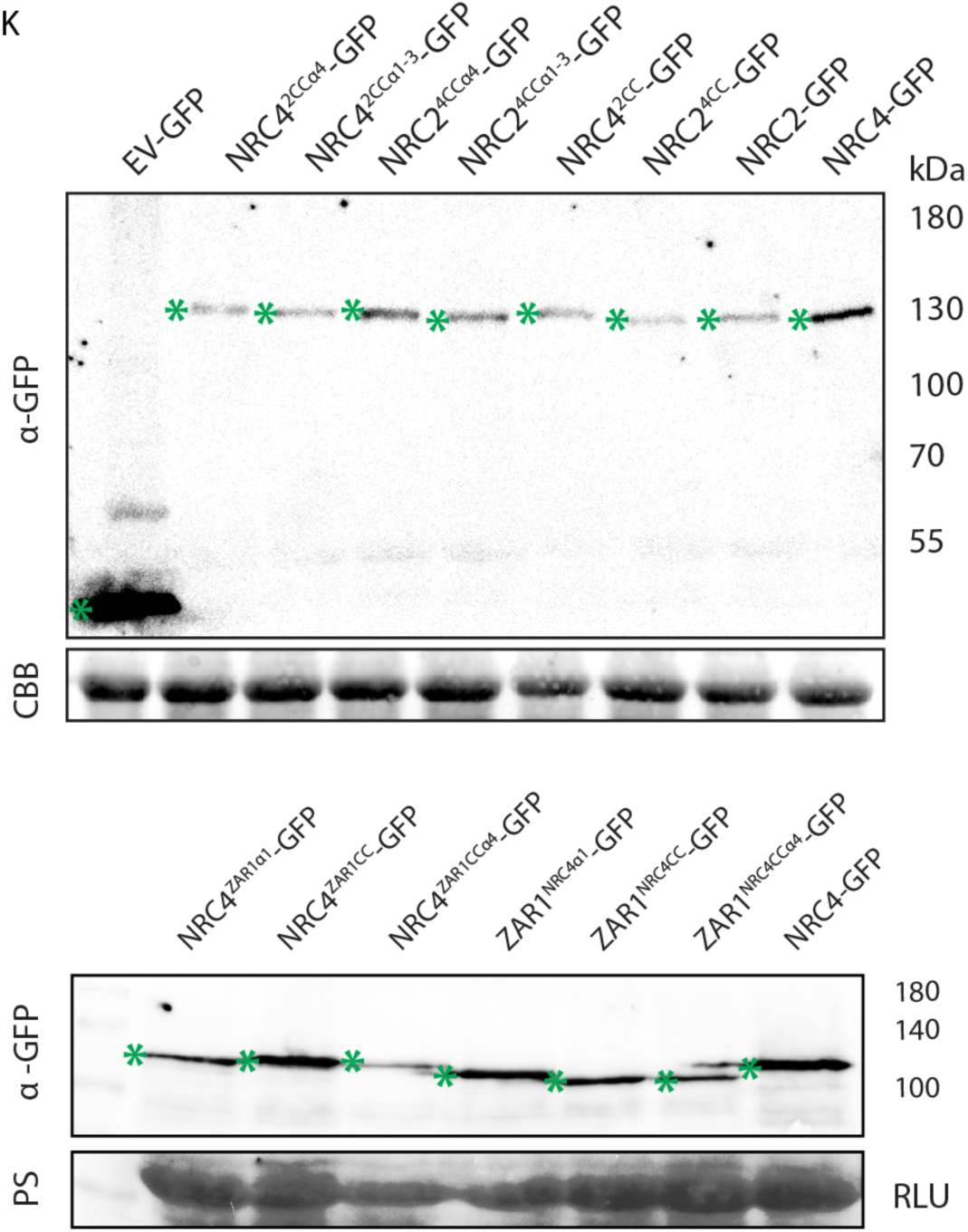
RFP-Rem1.3, BFP-EV and Merge channels for Figure 4, western blots showing chimeras are expressed. Single-plane confocal micrographs of C-terminally tagged GFP constructs of NRC4 truncates, NRC4-NRC2 chimeras or NRC4-ZAR1 chimeras, co-expressed in nrc4a/b plants with RFP-Rem1.3 and BFP-EV, with P19 for all but 4CC-GFP due to its high expression. Leaves were infected 3 hours post infiltration and imaged 3 days later. Arrowheads indicate haustoria. (A) NRC4ΔCC-GFP localises to the cytoplasm throughout the cell, including around the haustorium, but does not accumulate to the EHM; it also forms some (motile) puncta in the cytoplasm. (B) 4CC-GFP localises to the cytoplasm throughout the cell, including around the haustorium, but does not accumulate to the EHM. (C) NRC4^2CC^-GFP accumulates at the EHM. (D) NRC4^Z1CC^-GFP accumulates at the EHM, particularly around the haustorium neck. (E) ZAR1^4CC^-GFP localises to the cytoplasm throughout the cell, including around the haustorium, but does not accumulate to the EHM. (F) NRC2^4CC^-GFP accumulates at the EHM. (G) NRC2^4CCα4^-GFP accumulates at the EHM and forms filaments like NRC2, but less frequently. (H) NRC2^4CCα1-3^-GFP localises to the cytoplasm throughout the cell, including around the haustorium, shows some EHM labelling, but does not accumulate to the EHM. (I) NRC2-GFP localises to filaments dispersed throughout the cytoplasm, some of which associate with EHM, but NRC2 does not focally accumulate at the EHM. (J) NRC4-GFP accumulates at the EHM. (K) Western blotting shows expression of chimeras. Chimeras were expressed in nrc4a/b plants for 3 days with P19 to boost expression and 4 leaf disks were collected using #4 size cork borer RLU is rubisco large subunit stained by Coomassie Brilliant Blue (CBB) or Ponceau Stain (PS). Chimeras are all expected to be approximately 130 kilodaltons (kDa).

**Fig. S4.**
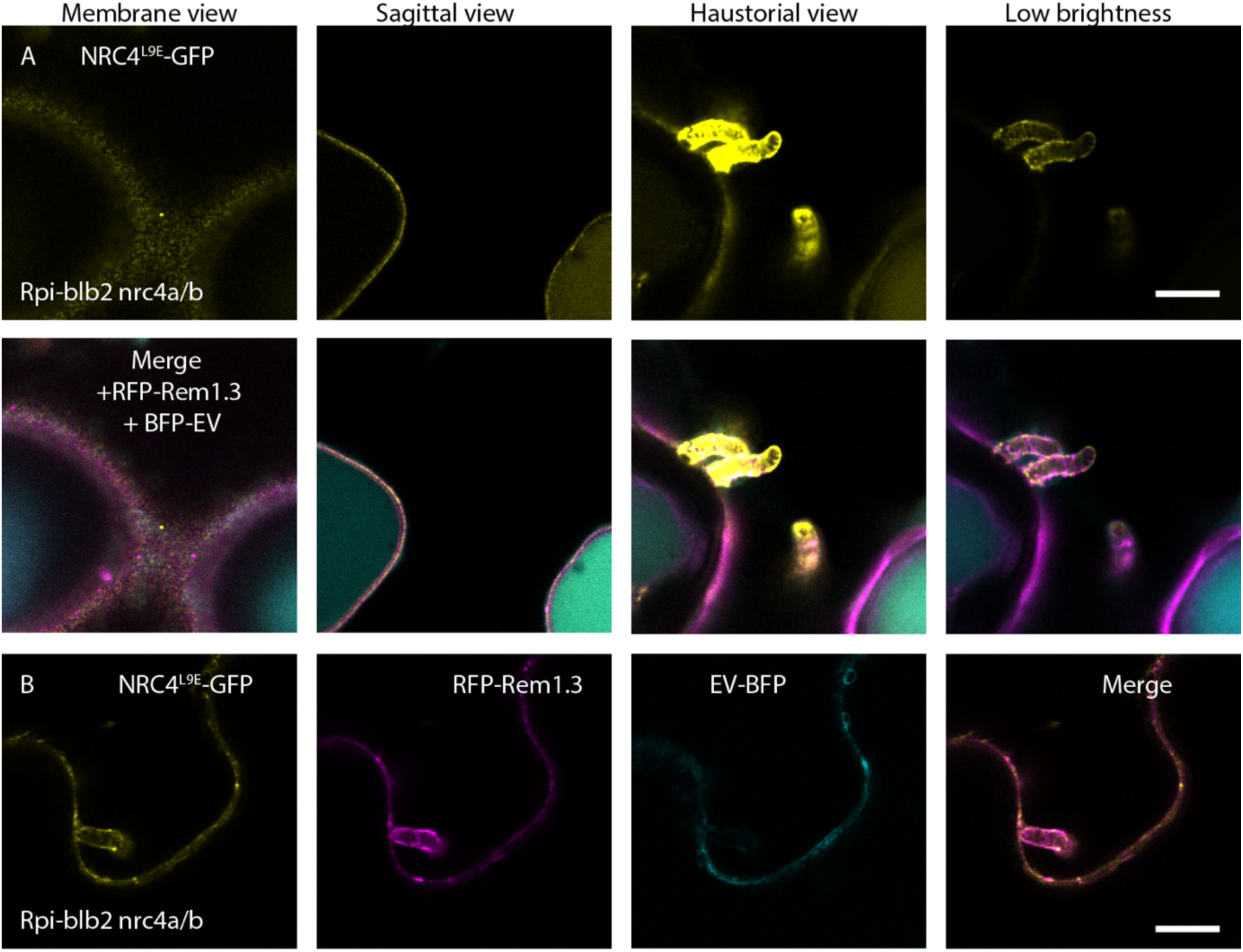
NRC4 activated at the EHM by Rpi-blb2 can form puncta at the EHM and plasma membrane. (A-B) Single-plane confocal micrographs showing the localisation of NRC4, with plasma membrane marker RFP-Rem1.3 and/or EV-BFP during infection with *P. infestans* in Rpi-blb2^nrc4a/b^ plants. All scale bars are 10 μm. (A) Example where NRC4^L9E^-GFP accumulates at the EHM but forms puncta on the plasma membrane in Rpi-blb2^nrc4a/b^ plants. Three Z-slices are shown to display the plasma membrane and EHM association. (B) Example where NRC4^L9E^-GFP does not focally accumulate but labels EHM and plasma membrane to the same degree and forms puncta associated with the EHM and plasma membrane in Rpi-blb2^nrc4a/b^ plants.

## Supplementary Movies

**Movie S1 (separate file): NRC4 accumulates around haustoria, EV-GFP remains cytoplasmic.** Z-stack of 14 slices. Transiently expressing EV-GFP (yellow) with NRC4-mKOk (magenta) during infection with *P. infestans.* Captured with Leica SP5.

**Movie S2 (separate file): NRC2 does not accumulate at the EHM but localises to filaments dispersed throughout the cytoplasm, but some of which associate with the EHM.** Z-stack of 18 slices. Transient expression of NRC2-GFP (yellow) co-expressed with EHM and plasma membrane marker RFP-Rem1.3 (magenta), during infection with *P. infestans*. Captured with Leica SP8.

**Movie S3 (separate file): NRC4 does not travel via RPW8.2 positive puncta implicated in RPW8.2 EHM accumulation.** Time lapse with frame interval 6 s, 18 frames. NRC4-mKOk (magenta) transiently co-expressed with RPW8.2-YFP (yellow) during infection with *P. infestans.* Captured with Leica SP5.

**Movie S4 (separate file): NRC4 associates to the plasma membrane, accumulates at the EHM and forms puncta associated with both when expressed in the presence of the sensor Rpi-blb2.** Z-stack of 16 slices. Transient expression of NRC4-L9E-GFP (yellow) with cytoplasmic marker EV-BFP (cyan) and plasma membrane and EHM marker RFP-Rem1.3 (magenta) during infection with *P. infestans.* Captured with Leica SP8.

## Notes

### Competing Interest Statement

SK receives funding from industry on NLR biology.

