## Supplementary material for "Dynamic accumulation of a helper NLR at the plant-pathogen interface underpins pathogen recognition": Dataset S1 Image Macro: Dataset S1 Chimera quantify macro.ijm

```
run("Duplicate...", "duplicate channels=1-3");
run("Median...", "radius=1");

macro "Multiple line Intensity Profile Plot Action Tool - C0g0 L1h43 L43hd C00g L1d6e L6eh4 Cg00 L1649 L49h8 Cg00 L1649 L49h8 C000 T3408l T6408i T9408n Tf408e L000h L0hhh C000 T3h07R T9h07O Thh07I"
{

run("Set Measurements...", "area mean min stack display redirect=None decimal=5");

//constants
colours = newArray("#4dff0000","#4d00ff00", "#4d0000ff","#4dff00ff","#4d00ffff","#4dffff00","#4dffc800","#4dffffff","#4d000000");
setTool("line");
lineWidth=1;
n=roiManager("count");


//get image info
fn=getTitle();
if (bitDepth()==24){
run("Make Composite");
}

getDimensions(ImageWidth, ImageHeight, ImageChannels, ImageSlices, ImageFrames);
getVoxelSize(px, py, pz, units);

if (ImageSlices>1 || ImageFrames>1){
	exit("This macro does not work on a Z or time series");
}


//get info
Dialog.create("Select Colour and Range");
	opt = newArray("Yes","No");
	if (n>0){
		Dialog.addChoice("Do you want to keep existing ROI's", opt, "Yes")
	}
	items = newArray("red","green","blue","magenta","cyan","yellow","orange","black","white");
	Dialog.addChoice("Colour", items, "green");

	Dialog.addChoice("Line Semi-Transparent ?", opt, "Yes")
	Dialog.addNumber("Line width", lineWidth);
	Dialog.addString("ROI Name", "Line", 12) 
Dialog.show();

if (n>0){
answer=Dialog.getChoice();
if (answer=="No"){
roiManager("Reset");	
}
}

col=Dialog.getChoice();
transparent=Dialog.getChoice();
lnwdth=Dialog.getNumber();
pre=Dialog.getString() ;

run("Line Width...", "line="+lnwdth);

colour=col;

//get transparent colour 
if (transparent=="Yes"){
for (ch=0;ch<9;ch++){
	if (col==items[ch]){
	colour=colours[ch];
}
}
}

//draw line and get data
waitForUser("Draw Profiles", "Draw lines and add to ROI manager (t) \n \n then select OK to continue");

n=roiManager("count");
if(n==0){
	exit("No ROI's")
	};
roiManager("deselect");
roiManager("Remove Channel Info");
roiManager("Remove Slice Info");
roiManager("Remove Frame Info");


//main routine
for (line=0; line<n; line++)
{
roiManager("select", line);
name=pre + " " + (line+1);
roiManager("Rename", name);
roiManager("Set Color", colour);
roiManager("Set Line Width", lnwdth);

}
roiManager("Deselect");


//start of analysis
//plots an intensity profile of a multi channel image

run("Clear Results");

bit=bitDepth();
yscalelim=pow(2, bit)-1;

//menu
Dialog.create("Plot size");
  Dialog.addMessage("Select Plot options: \n - if the colours appear wrong, you may need to re-set the LUT's\n - use the Change channel LUT's macro");
  range = newArray(" Set own limits ", " Full intensity range ", " Limit to upper signal "," Limit to signal range ");
  Dialog.addRadioButtonGroup("Select intensity range (y-axis)", range, 1, 4, " Limit to signal range ");
  Dialog.addNumber("Low intensity value ", 0, 0, 6," " );
  Dialog.addNumber("High intensity value ", yscalelim, 0, 6," " );
  Dialog.addMessage("Enter dimensions of plot window");
  Dialog.addNumber("X dimension ", 550, 0, 4," pixels" );
  Dialog.addNumber("Y dimension ", 300, 0, 4," pixels" );
  Dialog.addMessage("Set line width");
  Dialog.addNumber("Line width ", 1, 0, 4," pixels" );
Dialog.show;

rangelimits = Dialog.getRadioButton();
lowy = Dialog.getNumber(); 
highy = Dialog.getNumber(); 
plotsizex = Dialog.getNumber(); 
plotsizey = Dialog.getNumber();
plotlinewidth = Dialog.getNumber();  


for (line=0;line<n;line++){
selectWindow(fn);
roiManager("select", line);

Channel=getProfile();
xscalelen=lengthOf(Channel);


//get y range
highlevel=0;
lowlevel=yscalelim;
for (i=1;i<=ImageChannels;i++){
if (ImageChannels>1){
Stack.setChannel(i);
}
Channel=getProfile();
Array.getStatistics(Channel, lowtemp, hightemp, mean, stdDev);

if (hightemp>highlevel){
	highlevel=hightemp;
}
if (lowtemp<lowlevel){
	lowlevel=lowtemp;
}
}
extend=(highlevel-lowlevel)*0.025;
highlevel=highlevel + extend;
lowlevel=lowlevel - extend;


//get x range
	Distance=newArray(xscalelen);
for (g=0;g<xscalelen;g++){
	Distance[g]=g*px;
setResult("Distance (" + units +")", g, Distance[g]);
}


//y range selection
if (rangelimits==" Full intensity range "){
	ymax=yscalelim;
	ymin=0;
}
if (rangelimits==" Set own limits "){
	ymax=highy;
	ymin=lowy;
}
if (rangelimits==" Limit to upper signal "){
	ymax=highlevel;
	ymin=0;
}
if (rangelimits==" Limit to signal range "){
	ymin=lowlevel;
	ymax=highlevel;
}


//main plot and results
        Plot.create(fn+ " Intensity Plot Line "+line+1, "Distance (" + units + ")", "Intensity");
        Plot.setLimits(0, xscalelen*px, ymin, ymax);
        Plot.setFrameSize(plotsizex, plotsizey);
        Plot.setLineWidth(plotlinewidth);
for (ch=0;ch<ImageChannels;ch++){
if (ImageChannels>1){
Stack.setChannel(ch+1);
}
col=lookup();
Channel=getProfile();
chname="Channel " + ch+1 +" Line "+line+1;
for (row=0; row<lengthOf(Channel); row++)
{
setResult(chname, row, Channel[row]);
}
		Plot.setColor(col);     
        Plot.add("line",Distance, Channel);
        Plot.addText("Channel " + ch+1, ch*0.2, 0);
        Plot.setColor("Black"); 
        Plot.addText("Line " + line+1, 0.8, 0);
}
Plot.show();
}

run("Images to Stack", "name=Stack title=Plot use");
rename(fn+" Intesnity Profile Plots");

IJ.renameResults(fn+" Data");

selectWindow(fn);
run("Stack to RGB");
rename("Temp RGB");
roiManager("Show all with labels");
run("Select None");
run("Flatten");
rename(fn+" overlay");
selectWindow("Temp RGB");
close();


//exit();
pos_data=newArray(n*ImageChannels);
GFP_int=newArray(n*ImageChannels);
i=0;
selectWindow(fn+" Data");
Dist=Table.getColumn("Distance (microns)");
	
for (line=1;line<=n;line++){
GFP=Table.getColumn("Channel 3 Line "+line);
for (ch=1;ch<=ImageChannels;ch++){
Channel_data=Table.getColumn("Channel "+ch+" Line "+line);
Dist_pos=Array.reverse(Array.rankPositions(Channel_data));
pos_data[i]=Dist[Dist_pos[0]];
GFP_int[i]=GFP[Dist_pos[0]];
i=i+1;
}
}

//Array.show(pos_data,GFP_int);
titles=newArray("BFP Peak Position","RFP Peak Position","GFP Peak Position");
titles2=newArray("BFP Peak GFP Intensity", "RFP Peak GFP Intensity", "GFP Peak Intensity" );
i=0;
Table.create(fn+" Channel Peak Location");
Table.setLocationAndSize(400, 300, 800, 300);
for (line=0;line<n;line++){
Table.set("Line", line, "Line "+line+1);
for (ch=1;ch<=ImageChannels;ch++){
Table.set(titles[ch-1], line, pos_data[i]);
Table.set(titles2[ch-1], line,GFP_int[i]);
i=i+1;
}
}
}
//colour lookuproiManager("Show all with labels");
function lookup(){
getLut(reds, greens, blues);
Array.getStatistics(reds, min, max, mean, stdDev);
re=max;
RED=toHex(re);
if (re<17){
RED=RED+"0";	
}
Array.getStatistics(greens, min, max, mean, stdDev);
gr=max;
GREEN=toHex(gr);
if (gr<17){
GREEN=GREEN+"0";	
}
Array.getStatistics(blues, min, max, mean, stdDev);
bl=max;
BLUE=toHex(bl);
if (bl<17){
BLUE=BLUE+"0";	
}
col="#"+RED+GREEN+BLUE;
if (col=="#ffffff"){
col="Black";
}
return col;
}
}
```
